# Fingerprints of brain disease: Connectome identifiability in cognitive decline and Alzheimer’s disease

**DOI:** 10.1101/2022.02.04.479112

**Authors:** Sara Stampacchia, Saina Asadi, Szymon Tomczyk, Federica Ribaldi, Max Scheffler, Karl-Olof Lövblad, Michela Pievani, Aïda B. Fall, Maria Giulia Preti, Paul G. Unshuld, Dimitri Van De Ville, Olaf Blanke, Giovanni B. Frisoni, Valentina Garibotto, Enrico Amico, the Alzheimer’s Disease Neuroimaging Initiative

**Affiliations:** Department of Radiology and Medical Informatics (DRIM), Faculty of Medicine, University of Geneva, Geneva, Switzerland; Neuro-X Institute, École Polytechnique Fédérale de Lausanne, Geneva, Switzerland; Laboratory of Neuroimaging of Aging (LANVIE), University of Geneva, Geneva, Switzerland; Division of Radiology, Geneva University Hospitals, Geneva, Switzerland; Neurodiagnostic and Neurointerventional Division, Geneva University Hospitals, Switzerland; Lab of Alzheimer’s Neuroimaging & Epidemiology, IRCCS Istituto Centro San Giovanni di Dio Fatebenefratelli, Brescia, Italy; Faculty of Medicine, University of Geneva, Geneva, Switzerland; CIBM Center for Biomedical Imaging, Switzerland; Department of Psychiatry, Geneva University Hospitals, Geneva, Switzerland; Geneva Memory Center, Department of Rehabilitation and Geriatrics, Geneva University Hospitals, Switzerland; Division of Nuclear Medicine and Molecular Imaging, Geneva University Hospitals, Geneva, Switzerland; CIBM Center for Biomedical Imaging, Geneva, Switzerland

## Abstract

In analogy to the friction ridges of a human finger, the functional connectivity patterns of the human brain can be used to identify a given individual from a population. In other words, functional connectivity patterns constitute a marker of human identity, or a ‘*brain fingerprint*’. Yet remarkably, very little is known about whether brain fingerprints are preserved in brain ageing and in the presence of cognitive decline due to Alzheimer’s disease (AD). Using fMRI data from two independent datasets of healthy and pathologically ageing subjects, here we show that individual functional connectivity profiles remain unique and highly heterogeneous across early and late stages of cognitive decline due to AD. Yet, the patterns of functional connectivity making subjects identifiable, *change* across health and disease, revealing a functional reconfiguration of the brain fingerprint. We observed a fingerprint change towards between-functional system connections when transitioning from healthy to dementia, and to lower-order cognitive functions in the earliest stages of the disease. These findings show that functional connectivity carries important individualised information to evaluate regional and network dysfunction in cognitive impairment and highlight the importance of switching the focus from group differences to individual variability when studying functional alterations in AD. The present data establish the foundation for clinical fingerprinting of brain diseases by showing that functional connectivity profiles maintain their uniqueness, yet go through functional reconfiguration, during cognitive decline. These results pave the way for a more personalised understanding of functional alterations during cognitive decline, moving towards brain fingerprinting in personalised medicine and treatment optimization during cognitive decline.

## Introduction

The sharp increase of publicly available neuroimaging datasets ^1–3^ in the last few years has provided an ideal benchmark for mapping functional and structural connections in the human brain. At the same time, quantitative analysis of connectivity patterns based on network science has become more commonly used to study the brain as a network ^4,5^, giving rise to the scientific field of *Brain Connectomics* ^6^. Seminal work in this research area ^7,8^ has paved the way towards the new promising avenue of detecting individual differences through brain connectivity features. These studies showed that an individual’s functional connectivity patterns estimated from functional magnetic resonance imaging (fMRI) data, also known as functional connectomes (FCs), can constitute a marker of human uniqueness, as they can be used to identify a given individual in a set of functional connectivity profiles from a population ^7^. Given the analogy to well-known properties of the papillary ridges of the human finger, the field has taken the name of ‘brain fingerprinting’ and, since then, the extraction of “fingerprints” from human brain connectivity data has become a new frontier in neuroscience.

The excitement produced by the discovery that brain fingerprints can be extracted from matrices summarising human brain activity, either during rest or when performing a task, is unsurprising, for several reasons. Firstly, it confirms that studying the brain as a network can provide useful tools to get insights into the individual features that distinguish our brains one from the other; and second, it has been shown that brain fingerprints relate to behavioural and demographic scores ^9,10^. Accordingly, efforts have been made to implement ways of maximising and denoising fingerprints from brain data ^11–13^. These findings incentivized human neuroimaging studies to advance from inferences at the population level to the single-subject level, and allowed the field to move towards an individualised prediction of cognition and behaviour from brain connectomes ^14–17^. The next natural step is to explore whether this property of the human brain is maintained during disease. Despite promising preliminary findings towards this direction^18,19^, it is to date unclear to what extent FC-fingerprints could be used for mapping disease from human brain data. In fact, the vast majority of studies on brain fingerprinting have focused on healthy individuals, leaving the field of brain fingerprinting within the context of brain diseases as largely, if not completely, unexplored.

Cognitive decline and dementia are the final consequence of a series of pathological brain events. In the case of Alzheimer’s Disease (AD), which is the most common cause of dementia, these events involve the accumulation of toxic proteins such as β-amyloid and hyperphosphorylated tau between and within neurons, leading to neuronal death and ultimately causing damage to the wider structural and functional architecture ^20^. In line with this, AD is often referred to as a ‘disconnection syndrome’ ^21^ and, numerous studies have focussed on connectivity alterations in AD ^22–24^. This extensive body of literature has demonstrated that AD is associated with loss of functional connectivity between brain regions and disruption of network organisation ^25,26^. For instance, some studies have revealed that the observed loss of connectivity in certain circuits is often accompanied by hyperconnectivity in other brain regions/networks ^25–28^, and phases of hyperconnectivity may precede the onset of cognitive symptoms ^29–32^. Additionally, cognitive impairment in AD is associated with changes in *hub* regions ^33,34^, which play a crucial role in integrating information from domain-specific systems in the healthy brain ^4,35^. These *hub* regions often coincide with epicentres of β-amyloid accumulation ^33,34,36^, possibly due to their high metabolic needs ^34^. Moreover, AD patients show reduced nodal centrality in these *hub* regions like the medial temporal lobe and default mode network (DMN) ^37^. On the other hand, the strength of connectivity in tau epicentres predicts the topography of tau accumulation ^38,38–40^ and several groups are exploring the potential of exploiting functional connectivity as mean to predict future tau spread, with potential direct implications in clinical practice ^38,40^. Finally, graph theoretical measures indicate that AD is characterized by shorter path lengths between connected regions and an increased number of *hubs*, altering the global network organization ^26^. Overall, the literature suggests that during pathological aging, the brain undergoes both a loss and a reorganization of functional connectivity, and that these changes are closely tied to the underlying β-amyloid and tau pathophysiology.

However, to date, there is a lack of a functional connectivity alteration biomarker with an adequate level of specificity and sensitivity concerning to the various stages of AD ^23^, and this has hindered the routine use of fMRI in clinical practice ^30,41^. This is partly due to the intrinsic properties of connectivity from resting-state fMRI, a technique that is greatly influenced by factors affecting the obtained signal (heterogeneity in acquisition parameters, scanners characteristics, motion ^42^), by differences in the chosen signal-processing approaches ^30^, but also by the fact that existing studies focused on group averages, overlooking heterogeneity among individuals. The existence of significant inter-subject variability in terms of clinical manifestations is well-known in clinical practice, and this cognitive variability could be explained by individual features of functional organisation. Addressing the open research question of fingerprinting during brain disease could therefore open the door to individual characterization of cognitive decline from functional connectivity data and pave the way to a more widespread implementation in clinical settings. Investigating fingerprints of cognitive decline is also tightly connected to the concept of “precision medicine” ^43^, since it might not only provide insights on the individual trajectories of pathological brain ageing, but also allow for the surveillance of adapted personalised treatments during cognitive decline, advancing medicine in its quest for individualised biomarkers of neurodegeneration ^44^.

In this work, we investigated within-scan brain connectivity fingerprints using fMRI data collected from two independent datasets from individuals at different stages of cognitive decline. Within-scan fingerprinting, as introduced in ^12^, allows to uncover FC features leading to brain identification *within-session* and should be regarded as a *temporal stability investigation* of the resting-state functional brain architecture. We started by estimating functional connectome fingerprints of a clinically homogeneous and deeply characterised cohort of Cognitively Unimpaired amyloid β-negative individuals (CU Aβ-), amyloid β-positive patients with Mild Cognitive Impairment (MCI Aβ+) and amyloid β-positive patients with Dementia due to AD), during the first and second half of the fMRI session. We found that whole-brain functional connectivity patterns remained reliable across healthy and pathological brain ageing; in other words, it was possible to correctly identify a patient solely based on its functional connectome. Yet, significant differences in the spatial organisation of the brain fingerprint could be observed during cognitive decline. Notably, the functional connections that were the most reliable in healthy subjects disappeared during cognitive decline, leaving room for other stable connections, adjusted to the process of neurodegeneration. Furthermore, we looked at the distribution of connections with the highest fingerprint (i.e., temporally stable across time and allowing subjects identification) across cognitive decline, and we found a significant transition towards between-functional system connections, when going from healthy to dementia. Finally, this topological reconfiguration of brain fingerprints appeared to be associated to mostly high-order cognitive processes during healthy aging, and shifted to other functions during cognitive decline. Essentially, these findings demonstrate that functional connectivity profiles remain highly identifiable even during cognitive decline, thus providing crucial individualized information. This emphasizes the significance of harnessing this uniqueness by shifting the research focus from group differences to individual variability when investigating functional alterations in AD. Furthermore, we observed a functional reconfiguration of the brain fingerprint, providing valuable insights into specific functional connections and cognitive functions that could account for individual variability among individuals experiencing cognitive decline due to AD.

## Results

We investigated within-session brain fingerprints of neurodegeneration in two independent cohorts for a total of N=126 subjects (i.e., Geneva cohort: N=54 and ADNI: N=72) at different stages of cognitive decline: cognitively unimpaired Aβ-negative (CU Aβ-), with mild cognitive impairment Aβ-positive (MCI Aβ+), and with dementia due to AD (AD Aβ+). Our approach can be summarised in three steps: **(1)** We first estimated the functional connectomes (FCs) of each subject during the first and second halves volumes of fMRI acquisitions separately, (cf. Fig. 1A and see Methods for details). **(2)** We then estimated the degree of within-session brain identification or “brain fingerprint” at the *whole-brain level* for each group separately, through a mathematical object called identifiability matrix ^12^ (cf. Fig 1B). The identifiability matrix provides two useful metrics for brain fingerprinting: the degree of similarity of each subject with itself (*ISelf,* diagonal elements, Fig. 1B) as opposed to others (*IOthers*, off-diagonal elements Fig. 1B), and the degree of group fingerprint, conceptualised as the extent to which subjects were more similar to themselves than others (*IDiff*, see Methods). We also estimated the *Success-rate* of the identification procedure, as originally proposed by ^7^. **(3)** We further explored the *spatial specificity* of brain fingerprints by estimating the degree of distinctiveness of each FC-edge at the individual level, using intraclass correlation (ICC, see Methods and Fig. 1C). Edge-wise fingerprint was then computed at the networks (Fig. 1C) and hubs level (Fig. 1D).

**Figure 1.**
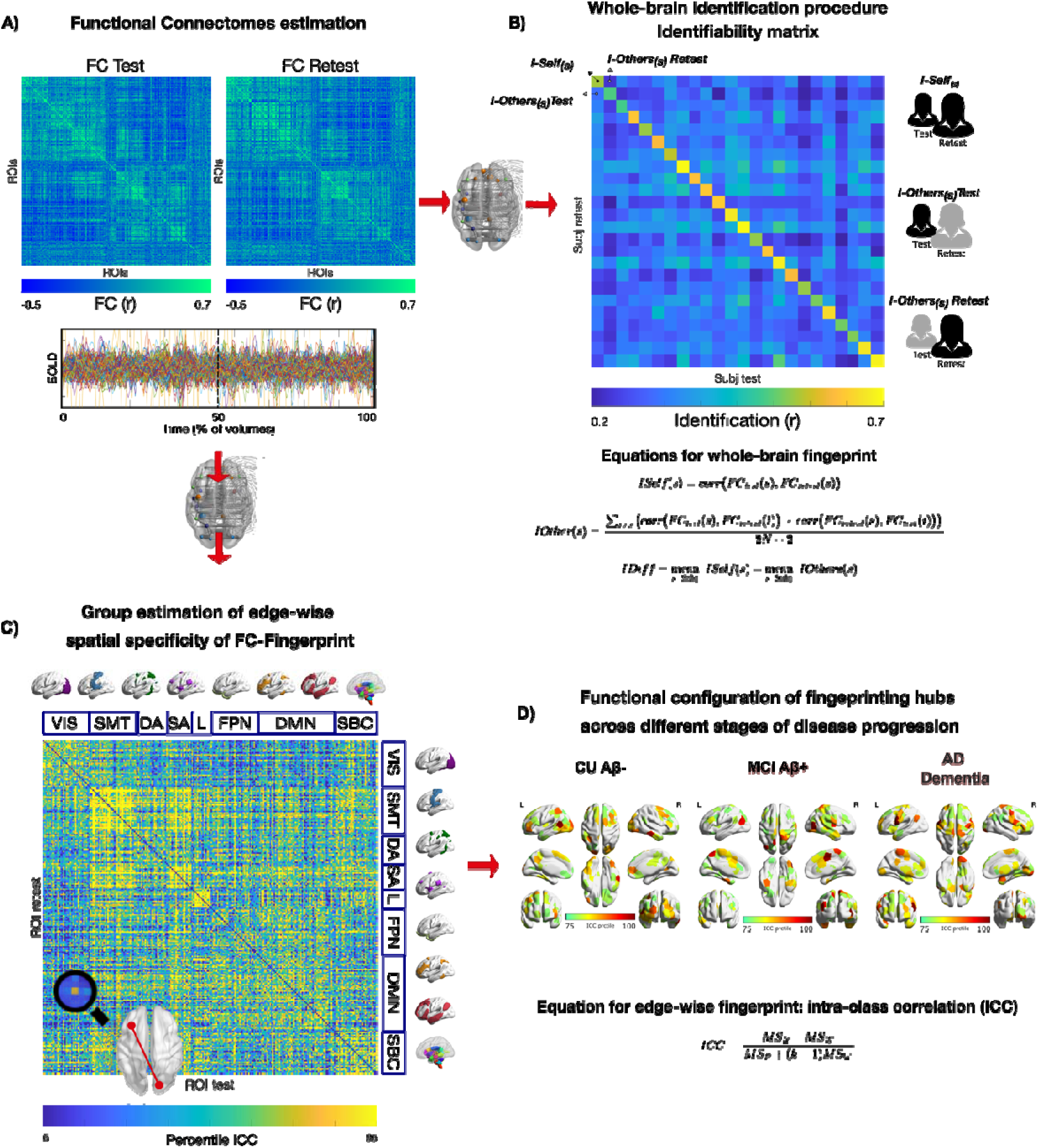
Exploring FC fingerprints of neurodegeneration, schematics of the approach. We estimated the FC-fingerprint in Cognitively Unimpaired Aβ-negative (CU Aβ-), Mild Cognitive Impairment Aβ-positive (MCI Aβ+) and Dementia Aβ-positive (AD Dementia) patients across two independent cohorts (Geneva, ADNI). Within-session fingerprint was estimated at the *whole brain level* as the degree of similarity between functional connectivity at test (FC Test, first 50% volumes) vs. retest (FC Retest, second 50%volumes, see Methods for details) **(A)** and summarised in a mathematical object called identifiability matrix **(B)** ^12^. Identifiability matrix shows within-subjects similarity (*ISelf*, elements in diagonal) and between-subjects similarity (*IOthers*, off-diagonal elements) across each group/cohort. Where *ISelf>IOthers* the identification procedure is successful. We also estimated *IDiff*, an estimation of the group-level whole-brain fingerprint as the distance between *ISelf* and *IOthers* (see methods). **C)** Spatial specificity of FC-fingerprint was estimated for each group/cohort using intraclass correlation (ICC), quantifying the fingerprint for each brain-edge (connection). ICC matrix is ordered according to the seven cortical resting state networks as proposed by ^45^. **D)** Fingerprinting hubs were computed as nodal strength of ICC matrix for each group. VIS=visual network; SMT=somatomotor network; DA=dorsal-attention network; SA=salience network; L=limbic network; FPN=fronto-parietal network; DMN=default-mode network; SBC=subcortical regions.

### Whole-brain within-sessions brain fingerprint during cognitive decline

In these first analyses, we investigated the degree of identification at the *whole-brain level*. We found that the *Success-rate* of the identification procedure was 100% in both the Geneva and ADNI cohorts. In other words, in each group each individual showed significantly higher similarity with themselves (*ISelf*), as opposed to others (*IOthers*) (Fig. 2B and 2D, Geneva and ADNI p<.0001), and *IDiff* was high in the three groups. We also found that test-retest reliability (*ISelf*) was high in the three groups and in both cohorts, with no significant differences across groups after controlling for nuisance variables, i.e., age, sex, years of education (YoE) and absolute difference between motion (FD) at test vs. retest [ANOVA with 5000 permutations to control for sample size differences; Geneva: F(2)=0.08, p=.918; CU Aβ-: M(SD)=0.60(0.07); MCI Aβ+: M(SD)=0.60(0.10); AD Dementia: M(SD)=0.60(0.07); cf. Fig. 2A; ADNI: F(2)=0.95, p=.389; CU Aβ-: M(SD)=0.73(0.07); MCI Aβ+: M(SD)=0.60(0.08); AD Dementia: M(SD)=0.71(0.08); cf. Fig. 2C]. See Fig. S1 for boxplots of *ISelf* across groups and Table 1A for full statistics for one-way ANOVAs. We tested differences in between-subjects similarity (*IOthers*) after controlling for nuisance variables (i.e., age, sex, YoE, average motion (FD) and scanner type, the latter for the ADNI dataset only). We found a main effect of group [ANOVA with 5000 permutations to control for sample size differences; Geneva: F(2)=15.3, p<.001; ADNI: F(2)=5, p=.013], revealing reduced between subjects similarity in MCI Aβ+ patients relative to CU Aβ-in the Geneva cohort [Bonferroni adjusted; MCI Aβ+ vs. CU Aβ-: p<.001] and in AD Dementia patients relative to CU Aβ-in ADNI [Bonferroni adjusted; ADNI: AD Dementia vs. CU Aβ-: p=.006]. See Fig. S1 for boxplots of *IOthers* across groups and Table S1B and S1C for full statistics for one-way ANOVAs and post-hoc pairwise comparisons. Finally, permutation testing showed that *IDiff* and *Success-rate* were different from null distributions at p <.0001 in all groups and in both cohorts (Fig. 2A, 2C). In sum, these data show that it is possible to correctly identify an individual with significantly greater accuracy relative to surrogate null models ^46^ (see Methods for details), independently from the clinical status, and solely based on the patterns of brain activity within-scan. We note that the identifiability results were replicated in two independent cohorts, using two different pre-processing pipelines.

**Figure 2.**
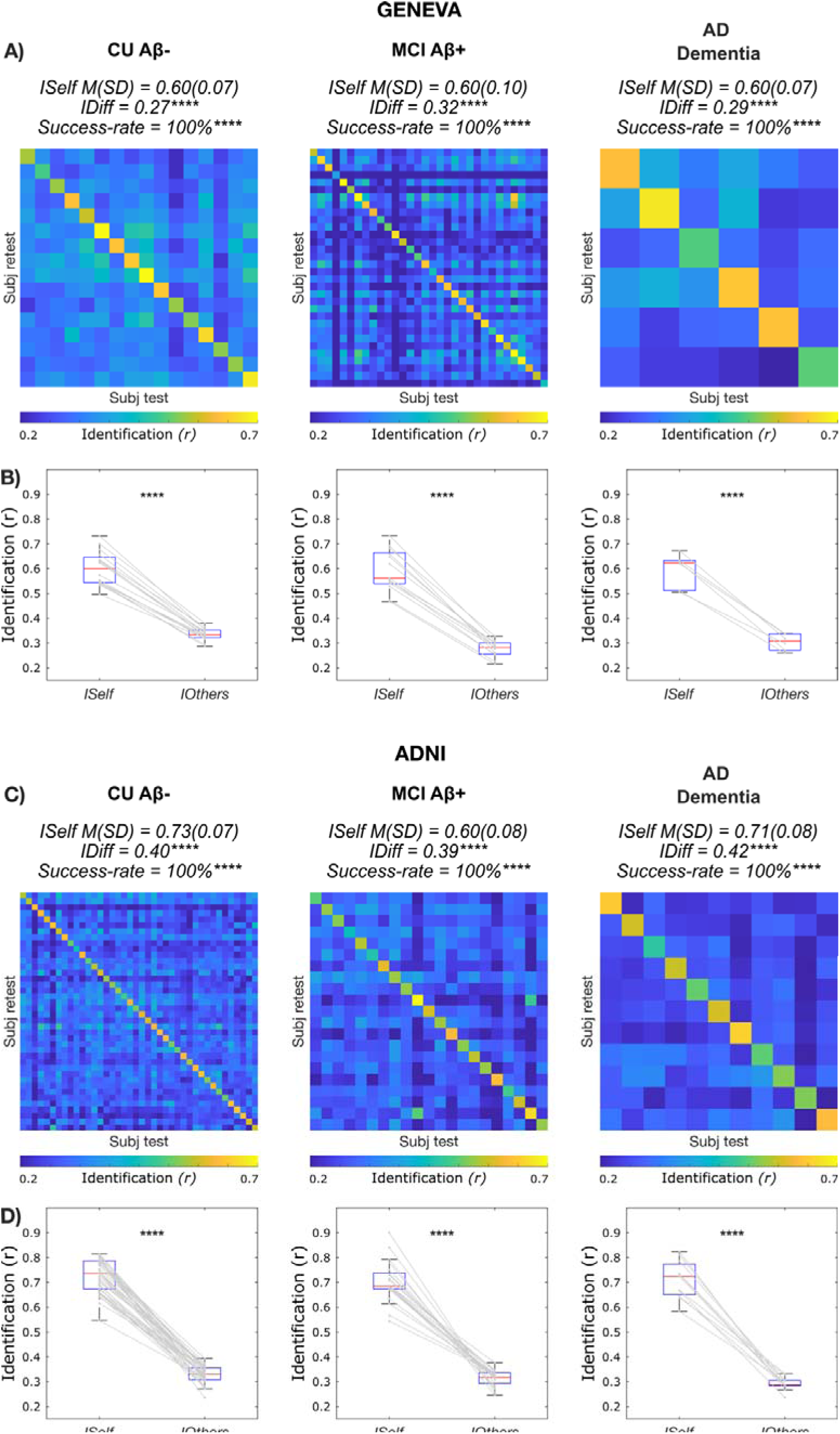
Functional Connectivity fingerprints during cognitive decline. **A-C)** Identifiability matrices show within-(*ISelf*) and between-subjects (*IOthers*) test-retest reliability as Pearson correlation coefficient in CU Aβ-, MCI Aβ+ and AD Dementia, for the two independent cohorts investigated (Geneva and ADNI, see Methods for details). Individuals’ *ISelf* and *IOthers* are displayed, respectively, in the diagonal and off-diagonal elements of the matrix. The average *ISelf, IDiff* and *Success-rate* were similar in the three groups and *IDiff* and *Success-rate* significantly differed from random distributions. **B-D)** Boxplots shows that *ISelf* was significantly higher (paired t-test) than *IOthers* in all individual cases and in all groups, for both the Geneva and the ADNI datasets. **** p ≤ 0.0001.

**Table 1.** Demographics - Geneva and ADNI.

| **Geneva (N=54)** | **CU Aβ-** | **MCI Aβ+** | **AD dementia** | **p** | **p signif.** |
| --- | --- | --- | --- | --- | --- |
|  | N=16 | N=32 | N=6 |  |  |
| Sex assigned at birth (% female) | 50.0 | 63.5 | 66.7 | 0.363 | ns |
| AGE M(SD) | 71(5.9) | 74.3(5.3) | 72.2(8.6) | 0.274 | ns |
| YoE M(SD) | 16.8 (3.4) | 12.8(4.6) | 11 (3.4) | 0.004 | ** |
| MMSE M(SD) | 28.6 (1.1) | 25.6(2.9) | 18.5(3.6) | <0.001 | **** |
| Centiloid M(SD) | -2.8(6.1) | 83.1(30.4) | 83.8(21.3) | <0.001 | **** |
| **ADNI (N=72)** | **CU Aβ-** | **MCI Aβ+** | **AD dementia** | **p** | **p signif.** |
|  | N=40 | N=21 | N=11 |  |  |
| Sex assigned at birth (% female) | 57.5 | 61.9 | 27.3 | 0.654 | ns |
| AGE M(SD) | 73.1(6.8) | 74.8(11.2) | 74.1(9.1) | 0.848 | ns |
| YoE M(SD) | 16.7(2) | 21(15.5) | 15.3(2.9) | 0.045 | * |
| MMSE M(SD) | 29.2 (0.9) | 27.2(2.4) | 21.6(2.7) | <0.001 | **** |
| Centiloid M(SD) | 6.2(9.8) | 80.6(36.1) | 105(36.1) | <0.001 | **** |

*Legend:* CU=cognitively unimpaired; MCI=mild cognitive impairment; Aβ -=Amyloid- $\beta$ status negative; Aβ + =Amyloid- $\beta$ status positive; MMSE=Mini-Mental State Examination; M(SD)=mean (standard deviation); p-value was estimated using Chi-square test for sex, and one-way ANOVA or its non-parametric equivalent, i.e., Kruskal-Wallis test for the remaining variables. ns=not significant, i.e., $p>0.05$ ; \* $p\leq 0.05$ ; \*\* $p\leq 0.01$ ; \*\*\* $p\leq 0.001$ ; \*\*\*\* $p\leq 0.0001$ .

The reported identification rates were computed at the whole-brain level, hence giving no information on the functional edges most influential for identification. The ensuing aim was therefore to understand how these results related to the local properties of the individual functional connectomes.

### Spatial specificity of brain fingerprint during cognitive decline

To address this question, we assessed spatial specificity of brain fingerprints using edgewise intra-class correlation ^12^ (ICC). In both cohorts, we observed a spatial reconfiguration of the most identifiable edges as cognitive decline progressed (cf. Fig. 3A and Fig. 3B). In other words, connections with the highest ICC had a different topological distribution in the three groups. Although there were differences between the two cohorts, in consistency with the variability of FC-patterns, we also identified common features that were shared among the groups. To investigate these common features, we examined the overlap between the edges with ‘good’ levels of ICC (ICC>0.6) ^47^ across the two cohorts (cf. overlap ICC matrix in Fig. 3C). We observed a slight decrease in the number of edges with ‘good’ ICC in MCI Aβ+ relative to CU Aβ-, and a large increase in AD Dementia relative to both CU Aβ- and MCI Aβ+ (cf. ‘n edges’ in Fig. 3C). The lower count of edges with high ICC in the MCI Aβ+ group may reflect the incipient AD pathology, but also the fact that MCIs Aβ+, despite being biologically homogeneous, can present with distinct clinical subtypes (e.g., amnestic vs non-amnestic) ^48^, with some level of uncertainty in disease trajectory (i.e., not all MCI Aβ+ convert to dementia ^49,50^). The first could lead to a reconfiguration of the functional architecture, the latter to less commonalities among individuals in terms of stable traits of functional connectivity, and both could explain to a reduced number of functional connections with stable patterns of connectivity across test and retest in all individuals. Conversely, the observed rise in AD Dementia patients could be attributed to a combination of greater clinical homogeneity and the advanced AD pathophysiology, which could impair brain ability to adapt and reconfigure, leading to an increased number of connections with stable test-retest connections. Moreover, we observed differences in spatial distribution across groups, suggesting a functional reconfiguration of the fingerprint from healthy to pathological ageing. Given these observations, we delved deeper into the spatial distribution of the edges with good ICC within each resting-state network.

**Figure 3.**
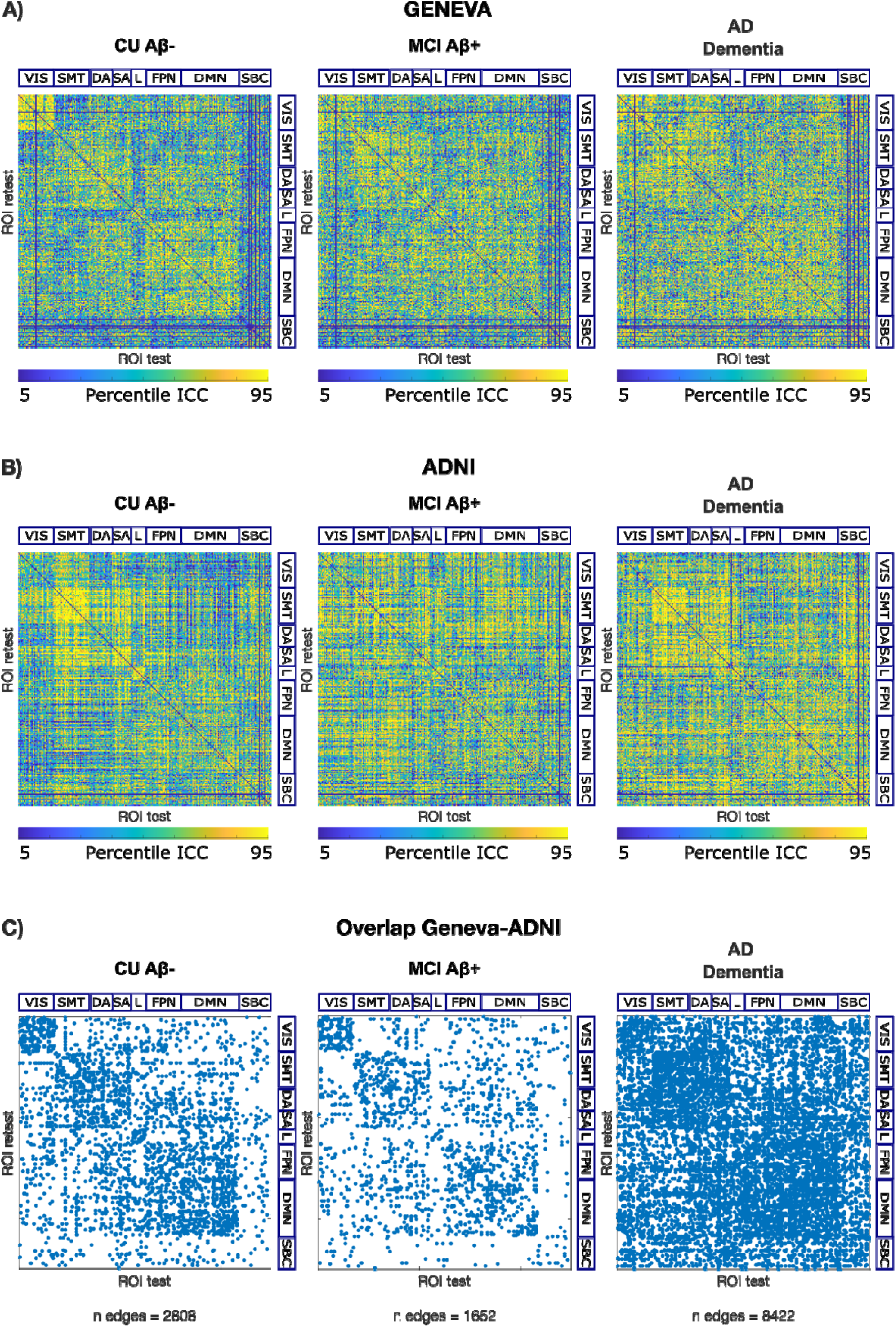
The connections with the highest fingerprint undergo spatial reconfiguration during cognitive decline. **A)** Geneva cohort; **B)** ADNI cohort; spatial specificity matrices of FC-fingerprints for each group as measured using edge-wise intra-class correlations (ICC). Here we display ICC values, computed using bootstrapping for randomly chosen N=10 subjects, across 1000 bootstrap runs, and then averaged within each group. We show edges with ICC between the 5^th^ and 95^th^ percentile. **C)** Overlap Geneva-ADNI; overlap across the two cohorts of spatial specificity matrices of FC-fingerprints. Matrices in A) and B) were binarized for ICC>0.6 which is considered a ‘good’ ICC score ^47^, and only overlapping edges are displayed. n edges=number of overlapping edges. Matrices in A), B) and C) are ordered according to seven cortical resting state networks (RSNs) as proposed by ^45^. VIS=visual network; SMT=somatomotor network; DA=dorsal-attention network; SA=salience network; L=limbic network; FPN=fronto-parietal network; DMN=default-mode network; SBC=subcortical regions.

### Brain fingerprint in resting-state functional networks during cognitive decline

We looked at the number of the edges in the binary overlap ICC matrix for each group (CU Aβ-, MCI Aβ+ and AD Dementia) in each network (within and between), relative to the total number of edges. Our results showed that MCI Aβ+ individuals had a slightly reduced fingerprint relative to CU Aβ-(i.e., slightly negative Ratio Disease/Health, R(net), with an average of −0.2 and −0.4 folds relative to CU Aβ-in within- and between-networks, respectively. In contrast, patients with AD Dementia showed a considerable increase in fingerprint (i.e., positive R(net)) across almost all functional networks, except for VIS-within and Limbic-within and between (see in Fig. 4A). The greatest increase was in the SBC network, including hippocampal and medial temporal regions, which are the earliest regions displaying atrophy and tau pathology in AD^51,52^. In addition, in between-networks there was an average increase of 2.9 folds, in particular for DMN, FPN and VIS. There was also higher fingerprint in within-networks, with an average increase of 2.2 folds. This shows that the increase in number of edges with highest ICC in AD Dementia, was widely distributed across the cortex but also especially driven by the regions with most long-lasting neuropathology, but also by between-network connections in key resting-state functional networks.

**Figure 4.**
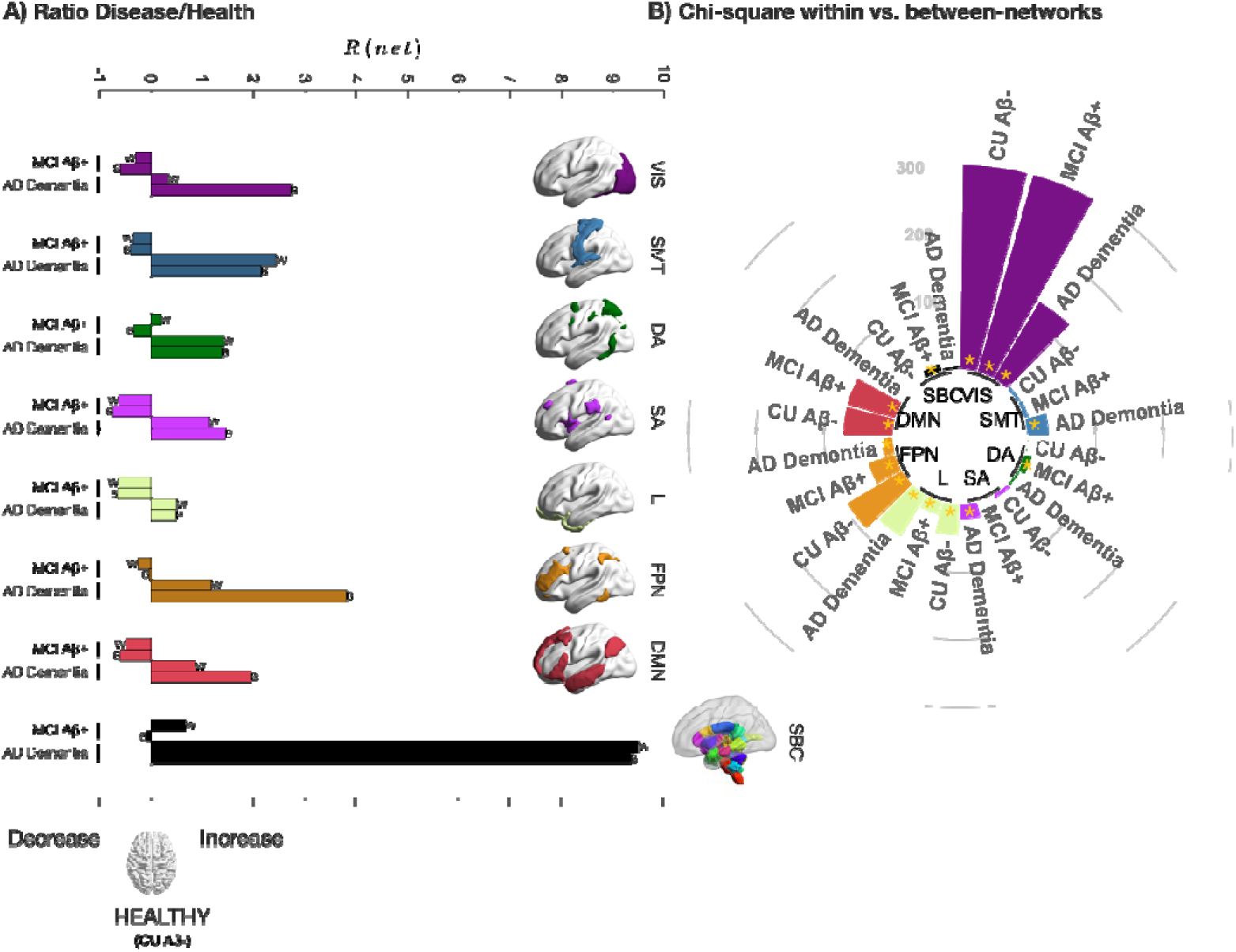
Distribution of brain fingerprint across resting-state functional networks. Distribution of the edges with highest ICC common to both cohorts (from ICC overlap binary matrix, cf. Fig. 3C) in within-networks and between-networks. **A)** Distance from healthy reference computed as ratio Disease/Health, cf. Methods). Positive/negative values denote increase/decrease in the percentages of edges, respectively. Note, across all networks, the overall increase in percentages of edges in AD Dementia patients and slight decrease in MCI Aβ+. **B)** Comparison of edges within vs. between networks, expressed as Chi-Square statistics. High value with * denote a significant (Bonferroni corrected) difference in the number of edges in within-networks vs. between-networks. With some exceptions (see main text and Supplementary Fig. 2), this reflects a higher percentage of edges in within relative to between-networks. Chi-Square for VIS in CU Aβ- and MCI Aβ+ was>800, but here we display Chi-Square ≤ 300 for visualisation purposes. W=within-networks; B=between-networks; ICC=intra-class correlation; CU Aβ-: Cognitively Unimpaired Aβ-negative; VIS=visual network; SMT=somatomotor network; DA=dorsal-attention network; SA=salience network; L=limbic network; FPN=fronto-parietal network; DMN=default-mode network; SBC=subcortical regions.

Next, we analysed the differences in within-network and between-networks fingerprints. We conducted a Chi-Square test to compare the number of edges within each network vs. the number of edges of that network with the others (i.e., between-network). The results showed that during healthy aging (CU Aβ-), in VIS, Limbic, FPN and DMN the proportion of edges with high fingerprint was significantly higher (Bonferroni corrected) in within-networks relative to between-networks (see Fig. 4B, see also Supplementary Fig. 2). Conversely, in both MCI Aβ+ and AD Dementia a notable reduction in Chi-Square statistics was observed in the FPN network. This indicates that, relative to controls, patients had higher fingerprint in the connections of the FPN with the rest of the brain. In Visual and DMN networks, a similar reduction in Chi-Square was observed, but only for AD Dementia, indicating higher number of edges in between-networks only in the advanced stages of the disease. For the Limbic network, we observed a reduction in Chi-Square for MCI Aβ+ and an increase during AD Dementia. This reveals a diminished discrepancy in edges with high fingerprint for between vs. within-network connections during the intermediate phases of the disease. Interestingly, during AD Dementia, the Limbic network exhibited a pattern similar to that of CU Aβ-, wherein an even higher proportion of edges with high fingerprint were found within the network, relative to between-network connections. Additionally, patients also showed increased fingerprint in between-network connections in other networks, where CU Aβ-did not show any notable within/between-networks proportion difference. For instance, MCI Aβ+ had increased fingerprint in between-networks connections in Subcortical and DA networks, while AD Dementia had similar increases in SMT network and significantly higher proportion in between-network connections for SA network. See also Supplementary Fig. 2, for the percentages of edges with high fingerprint in each network across groups.

In summary, these findings suggest that the distribution of edges with the highest fingerprint across resting-state functional networks undergoes changes during cognitive decline. Specifically, the distribution shifts towards more between-network connections in some key networks such as Visual, FPN and DMN, while in others, it remains relatively unchanged.

### Regional brain fingerprint during cognitive decline

Finally, we further explored the pattern of spatial reconfiguration as expressed by the ICC nodal strength of each brain region. Nodal strength was derived only from the edges that were significantly different from a permuted matrix randomly including subjects of the three groups (see Methods). We observed that regions contributing to the fingerprint (fingerprinting *hubs*) differed across groups and between cohorts (cf. Fig. S4), confirming that functional connectivity patterns are unique. Nevertheless, there was considerable overlap in the edges with the highest fingerprint across the two cohorts (Fig. 5).

**Figure 5.**
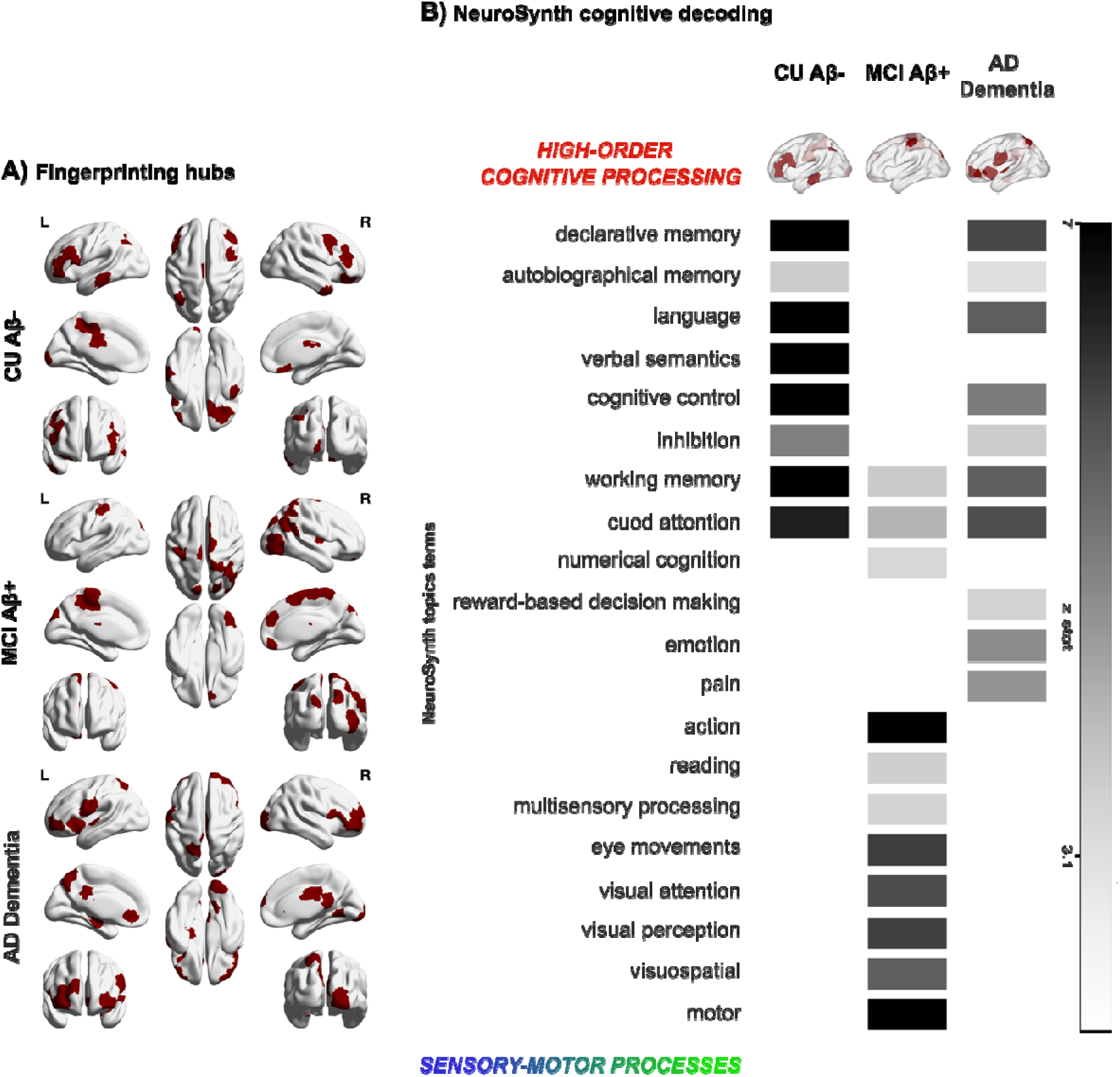
Fingerprinting *hubs* of cognitive decline and its association with behaviour. **A)** Brain fingerprint maps showing the top 25% brain nodes overlapping across the two cohorts. **B)** The Neurosynth meta-analysis of the brain fingerprints maps across cognitive decline shows a spectrum of association with higher order processes during healthy aging (CU Aβ-), towards lower-order motor-sensory processing during MCI Aβ+ and back to higher order, as well as affective and social processing, during AD Dementia. Brain fingerprints were linked with memory processes both in healthy aging and during cognitive decline, with different regions driving this association.

At last, we wanted to link brain fingerprints during the different stages of cognitive decline with behaviour. After deriving the group-specific nodes with the highest ICC values that were common across the two independent cohorts (cf. Methods), we applied a Neurosynth meta-analysis based on 50 topic terms onto the brain fingerprint of each group, similarly to previous work ^53,54^. We found that during healthy aging (CU Aβ-), brain fingerprints were associated with higher order processes such as long-term memory, language, semantics and executive functions, that rely on the integration of complex mental representations. On the other hand, in the initial phases of the disease and the incipient cognitive decline (MCI Aβ+) we observed a shift towards low-order sensory and motor processes, but also executive functions (‘working memory’, ‘cued attention’ and ‘numerical cognition’), ‘reading’ and ‘actions’. These regions, namely those involved in executive, visual, and motor processes, are progressively impacted in the later stages of AD. This pattern aligns with the ‘cascading network failure’ model, which posits a transition in connectivity from higher-order to lower-order circuits. This shift may represent an initial compensatory mechanism in the early stages of the disease, which eventually falters as the disease progresses^55^.

In the late stage of the disease (AD Dementia) we observed a shift-back to high-order cognitive functions, resembling the pattern observed in healthy aging, but also to regions implicated in affective/social processing including ‘pain’, ‘emotion’ and ‘reward-based decision making’. The involvement of regions modulating emotion and pain is consistent with clinical and imaging observations indicating that social-emotional functioning tends to remain relatively preserved in AD. This preservation aligns with hyperactivity observed in the SAL network, known for its role in detecting and integrating emotional and sensory stimuli^56^. In the context of the cascading network failure model, these circuits may represent a final opportunity for the brain to ‘compensate’ for AD pathology. On the contrary, highest fingerprints in regions implicated in higher order processes, may denote maladaptive stability in regions where the functional reconfiguration is no more possible. Note that brain fingerprints were linked with memory processes both in healthy aging and during AD Dementia (cf. Fig.5B), although the regions driving this association differed in the 3 groups (cf. Fig. 5A).

## Discussion

The pressing quest of neuroscience is to deepen the understanding of the intricate links between brain and cognition, behaviour, and dysfunction. Brain fingerprinting holds the potential to provide a crucial contribution to this endeavour, building upon its capacity to derive individual inferences from functional connectivity profiles. Seminal work ^7,12^ showed that functional connectivity profiles at rest are highly heterogenous in healthy individuals, especially in regions devoted to higher-order cognitive functions, such as such as Frontoparietal (FPN) and DMN, and this heterogeneity may reflect individual variability in cognition and behaviour. However, very little is known about how this property of the human brain may change as a consequence of cognitive decline and neurodegeneration. In this work, we aimed to provide an answer to the fundamental question of how human brain fingerprints changes during normal and pathological brain ageing due to Alzheimer’s Disease. To do so, we evaluated the within-session identification properties of functional connectomes during healthy aging, MCI and Dementia due to AD (i.e., MCI Aβ+ and AD Dementia) in two independent cohorts using fMRI data from N=126 individuals, and along two main lines: i) we investigated whether the within-session identification properties were maintained at the whole-brain level across the different clinical groups, ii) we explored the spatial configuration of brain fingerprints along the continuum between healthy and pathological brain aging due to AD (Fig. 1).

Firstly, our study revealed that individuals can be accurately identified based solely on the patterns of brain activity, regardless of their clinical status (Fig. 2). Remarkably, individuals remained highly consistent in their brain connectivity across test and retest sessions (*ISelf*), independent of their clinical condition. Additionally, patients exhibited a high degree of distinguishability from one another (*IOthers*), sometimes even more so than healthy elderly individuals among themselves (Fig. S1B). This finding suggests that disease can enhance the distinctiveness of individuals based on their functional connectivity. We note that these results were observed in two independent datasets (Geneva cohort, ADNI) with different acquisition parameters, signal-to-noise ratio and pre-processing pipelines, supporting the robustness of our results. These findings carry significant implications, as they reveal that functional connectomes remain highly heterogenous *also* during cognitive decline due AD and that they contain important individualised information. This highlights the importance to take into consideration, and make use of, this rich inter-individual variability to fully capture the complexity of functional alterations associated with cognitive decline and AD. Currently, the literature lacks consensus on the functional connectivity alterations that occur during different stages and causes of cognitive decline ^23^, and this has hindered the use of fMRI as a clinically-relevant tool ^30,41^. Previous studies have primarily focused on group averages, overlooking the heterogeneity among individuals and potentially disregarding essential information embedded within individual variability. For instance, evidence suggests that the distribution of pathological tau aggregates in the brain is linked to the functional connectivity architecture ^38–40,57^. If it is known that the spatial progression of tau aggregates in the cerebral cortex mirrors the severity of symptoms ^58,59^ and follows various deposition trajectories ^60^, being able to predict an individual’s trajectory would be crucial for the integration of disease progression profiles into clinical practice, and could also significantly impact research on disease-modifying therapies for AD ^61^ aiming to reduce pathological proteins accumulation and spread. Our findings carry important implications for this field of research, as they suggest that leveraging the individual characteristics of functional connectomes is fundamental to elucidating the heterogeneity in the patterns of tau spread and, therefore, disease progression.

When examining the topological distribution of connections underlying this uniqueness, we observed a spatial reconfiguration of regions with the highest fingerprint during different stages of Alzheimer’s Disease. Consistent with previous research utilizing magnetoencephalography-based functional connectomes ^18^, we identified a global decrease in identification between healthy elderly individuals and those with MCI. Additionally, and extending this work, we reported for the first time a significant increase in the number and sparsity of these temporally stable connections in patients with AD Dementia (Fig. 3C and Fig. 4A). In MCI Aβ+, the Alzheimer’s disease pathology is at its early stages, and it is conceivable that this can lead to functional connectivity reconfiguration and readaptation, resulting in a reduced number of connections with stable patterns of connectivity among individuals. In our study, we included only MCI Aβ+, which is a biologically homogeneous subtype with a relatively predictable clinical trajectory ^49,50^. However, not all MCI Aβ+ convert to Dementia and can present with distinct clinical subtypes (e.g., amnestic vs non-amnestic). Thus, we cannot exclude that the reduced number of regions contributing to the fingerprint may also reflect this clinical heterogeneity. Conversely, Dementia due to Alzheimer’s disease (AD) represents a more well-defined clinicopathological entity ^41,62^, and the advanced stage of the disease facilitate, in most cases, clinical exclusion of alternative aetiologies, i.e., differential diagnosis, leading to a higher clinical homogeneity between patients. In addition, the AD pathophysiology is advanced in these patients and the functional reconfiguration is likely to have reached its plateau, and this could result in higher number of connections remaining “unhealthily” stable across time among individuals. Furthermore, previous work has showed that patients with Dementia spend more time in sparse connectivity configurations ^63^, which may explain the greater number of sparse between-networks connections exhibiting high stability over time. Our results not only reveal increased stability across time in AD Dementia but also highlight between-subject heterogeneity in terms of connectivity strength, both of which are captured by our fingerprint metric, i.e., ICC ^12,64^. The heightened stability observed in AD Dementia patients is an unfavourable hallmark, potentially indicating that the brain is no longer flexible and dynamic. In contrast, the diminished temporal stability in MCI may denote an attempt to counteract pathological changes by enhancing dynamic interactions between neural circuits.

Next, we observed that during healthy aging, in the visual (VIS), fronto-parietal (FPN), default-mode (DMN) and limbic (L) networks, edges with the highest fingerprint were mostly within-networks, while they appeared increasingly in between-networks connections as cognitive decline started (MCI Aβ+) and progressed (AD Dementia) (Fig. 4B and Fig.3). Previous work consistently showed that the FPN and DMN networks are found amongst the networks that display the highest inter-subjects variability ^7,65,66^ and our results showed that the variability initially intrinsic to within-system connections gets distributed to connections between these networks and the rest of the brain, as cognitive decline progresses. Notably, our findings in AD Dementia patients showed that having increasing temporally stable and therefore differentiable links does not always carry a positive prognostic. One possible hypothesis would be that the neurodegenerative processes affect the healthy topological variability of functional connectivity patterns, creating unhealthy “hyperstability” in the functional connections between different functional systems, which then could hinder the capacity of the brain networks to hop ^67^ or reconfigure ^68^ between different dynamical states. This aligns with prior research demonstrating that variability in brain function is crucial for ensuring the brain’s optimal responsiveness to a dynamic environment, and that this characteristic diminishes with age^69–71^ and generally supports cognitive performance^70,71^.

Furthermore, previous works showed that the main drivers of the uniqueness of each individual functional connectome originates from brain areas responsible for higher-order cognitive processing during health ^7,12^. However, it was not known whether this changed in response to cognitive decline and Alzheimer’s Disease. In this work, we present strong evidence based on two independent cohorts, revealing how within-session brain fingerprints map onto different cognitive functions during healthy vs. pathological aging. Specifically, during healthy aging, brain fingerprints exhibit a range of associations with higher-order processes, resembling those observed in young, healthy individuals in between-sessions fingerprints. Conversely, in MCI Aβ+, brain fingerprints show a shift towards lower-order sensory-motor processing, as well as executive functions, ‘reading’ and ‘actions’. In the early stages of the disease, as amyloid pathology accumulates and affects connectivity within regions responsible for higher-order cognition, such as the DMN ^36^, it is plausible that these perturbations may lead to decreased regularity and stability in their connectivity patterns over time, resulting in a less distinctive ‘fingerprint’. In contrast, sensory-motor regions, typically unaffected by early amyloid pathology ^72^, may exhibit adaptive changes in their functional connectivity patterns as a compensatory mechanism. This is also in line with the ‘cascading network failure’ model proposed by Jones et al.^55^. Conversely, as Alzheimer’s Disease (AD Dementia) progresses to its advanced stages, we have observed a shift in fingerprint towards higher-order cognitive functions, encompassing affective and pain processing as well as decision making influenced by reward. This suggests that when the neuropathology is its advanced stage (including tau pathology and neurodegeneration), the functional reconfiguration processes in regions initially affected by amyloid pathology -primarily involved in higher-order cognition – tend to halt leading to a maladaptive stability, albeit with distinct levels of functional connectivity among individuals. This emphasizes the individualized nature of impairment in high-order functions like memory and executive functions.

Finally, our findings shed light on cognitive functions that are typically overlooked in memory-related conditions such as AD, e.g., somatomotor-processes and pain and affective functioning. Our data showed these to contribute significantly to the variability in functional connectivity across individuals in the early and latest stages of the disease, respectively. These findings may therefore have also implications for neuropsychological assessment and interventions, as they suggest that it may be important to broaden the focus to these overlooked cognitive functions in order to better tackle inter-individual variability. In essence, the observed gradient of association suggests a transition in the highly differentiable hubs from higher-order cognitive systems – associated to “healthy” abstract cognitive functions such as memory or semantics – to more “low-order” ancestral/sensory systems, during the early stages of the disease, and then back to high-order cognition to the late stages of the disease.

This study has some limitations. First, the impact of the choice of the brain atlas should be further verified. Second, it is known that connectivity measures are highly susceptible to artefacts arising from head motion and respiratory fluctuations ^73,74^, and these effects are even more pronounced in pathological conditions. However, in our datasets, we did not find high motion data points to significantly contribute to the differences in fingerprinting across groups. We observed no significant difference across groups in the percentage of censored volumes [p>=.066] or in the average FD [p>=.146], and differences in motion between test and retest volumes did not explain the variance of individuals’ test-retest similarity (i.e., ISelf, cf. Table S1A). Nonetheless, future work should analyse in depth the effect of motion at shorter time scales, where these artefacts can dominate. Third, in this study, we used two halves of the same scanning session to estimate identifiability, focusing more on the FC features leading to brain identification *within-session*. Obtaining test-retest sessions across different days in clinical cohorts poses significant challenges, and to the best of our knowledge, there are currently no publicly accessible large datasets of fMRI data collected across closely spaced time points (such as one week apart) in cognitively impaired cohorts, unlike with healthy cohorts (e.g., Human Connectome Project). One potential workaround to this issue is to cut the resting state time series in half, as originally proposed ^12^. Although this approach introduces the confound of looking at “within-session” fingerprinting, which could be influenced by the specific moment of scanning, it has the advantage of reducing scanner and acquisition noise, which are typically major confounding factors in connectome identification ^13^. Moreover, this method has been shown to yield similar identifiability results compared to data acquired across separate sessions (refer to Fig. S3 in ^12^ for a comparison on healthy subjects’ data from the Human Connectome Project). Nonetheless, it should be noted that within-scan fingerprinting in this work should be regarded more as a first *temporal stability investigation* of the resting-state functional brain network across cognitive decline. Future studies should aim to replicate our analysis using the “standard” between-sessions identification approach.

This work raises also important novel questions and indicates directions for upcoming research. For example, future works could build upon these findings by examining the temporal aspects of brain fingerprints in neurodegeneration. Recent studies have in fact demonstrated that, in healthy individuals, brain-fingerprints emerge at different time-scales, with for different networks/cognitive functions^75^, yet it is unknown whether this could change as a consequence of cognitive decline. It will also be important to characterize FC-fingerprints in a longitudinal dataset, i.e., to investigate whether the fingerprint is maintained after long time-gaps (e.g., years) and whether this changes in declining vs. stable individuals. Subsequent studies should also delve deeper into the relationship between atrophy, tau and amyloid accumulations, and changes in spatial patterns of brain fingerprints among patients. In our study, both cohorts were stratified based on amyloid status and stage of cognitive decline. Future investigations with larger subject numbers could explore differences in groups with stratification using comprehensive biomarker phenotyping, such as the presence of tau pathology, atrophy, and APOE genotyping.

### Conclusions

Functional connectivity patterns in the human brain exhibit remarkable distinctiveness, enabling the identification of healthy individuals within a population. In this work, we have discovered that this property of the human brain, known as the *brain-fingerprint*, is maintained during aging, and in the Alzheimer’s continuum, however differently configured. By investigating this topological reconfiguration and its link with cognition, we found that the heterogeneity amongst individuals was mostly driven by high-order cognition regions during healthy aging, with a shift towards lower-order sensory-motor regions in MCI Aβ+ and back to high-order cognitive and affective functions in AD Dementia. In essence, this work demonstrates that Alzheimer’s Disease significantly impacts the functional architecture of the human brain, albeit in remarkably unique ways for distinct individuals. These findings hold profound implications as they uncover functional connectivity as a personalized metric carrying substantial individualized information, highlighting the need to shift the research focus from group-averages to individual differences, and opening doors to various applications in personalized medicine. This work lays the foundation for clinical fingerprinting using functional magnetic resonance imaging, enabling a deeper understanding of cognitive decline at an individual level, with the potential to inspire novel approaches that leverage the distinct characteristics of functional connectivity, to enable personalized symptom monitoring, more accurate diagnosis, therapy surveillance and enhanced prediction.

## Materials and Methods

### Participants and demographics

Participants were included from two independent cohorts: the Geneva Memory Centre (Geneva University Hospitals, Geneva, Switzerland) and the Alzheimer’s Disease Neuroimaging Initiative (ADNI). Inclusion criteria were availability of (i) fMRI and T1-weighted scans, (ii) 18F-Florbetapir or 18F-Florbetaben amyloid-PET to derive amyloid β-status (iii) clinical and cognitive assessments and demographic data, and (iv) identical fMRI acquisition parameters (cf. ‘Image acquisition parameters’ sections). The exclusion criterion was the presence of any significant neurologic disease other than AD (cf. ‘Clinical assessment’ section). Subjects ranged from healthy ageing and Aβ-negative (cognitively unimpaired, CU Aβ-), to mild cognitive impairment Aβ-positive (MCI Aβ+), and Aβ-positive subject with dementia due to probable AD, AD dementia; cf. ‘Clinical assessment’ section for details about clinical and Aβ-status).

#### Geneva

N=58 subjects from the Geneva Memory Centre were included. Four subjects were excluded when motion-tagged volumes (see below) were>30%, leaving a total of N=54 remaining subjects for analyses. These included N=16 CU Aβ-, N=32 MCI CU Aβ+, and N= 6 patients with AD dementia (cf. Table 1 for all study-relevant covariates). Differences across groups in age, years of education and MMSE were tested using one-way ANOVA or its non-parametric equivalent, i.e., the Kruskal-Wallis test; Chi-square test was used for sex. There were no differences in age [p=.274] and sex [p=.363] across groups, while AD dementia and MCI Aβ+ were on average significantly less educated than healthy individuals [p=.004]. As expected, MMSE [p<.001] and Centiloid [p<.001] scores varied across groups [p<.001], revealing lower cognition and higher amyloid load in MCI Aβ+ and AD dementia patients relative to CU Aβ-.

#### ADNI

Data were obtained from the Alzheimer’s Disease Neuroimaging Initiative (ADNI) database (adni.loni.usc.edu). The ADNI was launched in 2003 as a public-private partnership, led by Principal Investigator Michael W. Weiner, MD. The primary goal of ADNI has been to test whether serial magnetic resonance imaging (MRI), positron emission tomography (PET), other biological markers, and clinical and neuropsychological assessment can be combined to measure the progression of mild cognitive impairment (MCI) and early Alzheimer’s disease (AD).

N=79 subjects from the ADNI database were included. Seven subjects were excluded when motion-tagged volumes (see below) were>30%, leaving a total of N=72 subjects for analyses. These were N=40 CU Aβ-, N=21 MCI Aβ+ and N=11 AD dementia patients (cf. Table 1 for all study-relevant covariates). Differences across groups in age, years of education and MMSE were tested using one-way ANOVA or its non-parametric equivalent, i.e., Kruskal-Wallis test; Chi-square test was used for sex. There were no differences in age [p=.848] and sex [p=.654] across groups, while AD dementia were on average significantly less educated than healthy individuals [p=.045]. As expected, MMSE [p<.001] and Centiloid [p<.001] scores varied across groups [p<.001], revealing lower cognition and higher amyloid load in MCI Aβ+ and AD dementia patients relative to CU Aβ-.

### Clinical assessment

Clinical status was established by expert neurologists of the Geneva Memory Centre (cf. for full details on the clinical assessment) for the Geneva cohort, and from ADNI collaborators for the ADNI cohort (cf. https://adni.loni.usc.edu/wp-content/themes/freshnews-dev-v2/documents/clinical/ADNI3_Protocol.pdf for full details on the clinical assessment). In brief, CU were individuals with or without subjective cognitive complaints and an absence of significant impairment in cognitive functions or activities of daily living (Geneva cohort: MMSE≥27, non-depressed; ADNI: MMSE≥27 and Clinical Dementia Rating (CDR)=0, non-depressed, cf. Table 1). MCI were subjects with objective evidence of cognitive impairment, cognitive concern reported by the patient and/or informant (family or close friend), and little or no functional impairment in daily living activities (Geneva: MMSE≥19) (ADNI: MMSE≥19, CDR=0.5). Individuals living with dementia were defined based on the same above criteria for MCI, but with impairment in the activities of daily living and fit the NINCDS/ADRDA criteria for probable AD (GENEVA: MMSE=12-20) (ADNI: MMSE=17–26, CDR≥0.5).

For both datasets the exclusion criterion was the presence of any other significant neurologic disease. These included: Parkinson’s disease, multi-infarct dementia, Huntington’s disease, normal pressure hydrocephalus, brain tumour, progressive supranuclear palsy, seizure disorder, subdural hematoma, multiple sclerosis, or history of significant head trauma followed by persistent neurologic deficits or known structural brain abnormalities (cf. https://adni.loni.usc.edu/wp-content/themes/freshnews-dev-v2/documents/clinical/ADNI3_Protocol.pdf).

### Amyloid-***β*** status

In the Geneva cohort, amyloid-β deposition was measured using ^18^F-florbetapir or ^18^F-flutemetamol PET, using standard imaging protocol, reconstructions and pre-processing pipelines, previously described in detail^76^. Given the use of two different amyloid-PET tracers, the standardized uptake value ratio (SUVr) was converted to a common scale, the Centiloid (CL) scale, a standardisation method proposed to harmonise the results obtained across tracers^77^. Aβ-status was determined in two ways: using the previously established cut-point (CL > 12^78^) and visually determined by an expert nuclear medicine physician (VG, >15 years of experience in the field) using visual assessment and standard operating procedures approved from the European Medicines Agency ^79,80^. In two discordant cases, where the CL was borderline, the visual assessment (positive) was preferred.

In ADNI, Aβ-status was determined using global amyloid-PET SUVR, derived after whole cerebellum normalisation of ^18^F-florbetapir or ^18^F-florbetaben PET, following pre-established reconstruction and pre-processing protocols (cf. https://adni.loni.usc.edu/methods/pet-analysis-method/pet-analysis/) and cut-points (global AV45 SUVR > 1.11; global FBB SUVR > 1.08)^81^. As in the Geneva cohort, to allow aggregation of data from the two tracers, the global amyloid-PET SUVR values were converted to the Centiloid scale^77^ and reported in Table 1.

### Image acquisition parameters

#### Geneva

Structural and functional data were acquired using a 3T Siemens Magnetom Skyra scanner (Siemens Healthineers, Erlangen, Germany) using a 64-channels phased-array head coil. Scans were performed within the radiology and neuroradiology division, Geneva University Hospitals, Geneva, Switzerland. An EPI-BOLD sequence was used to collect functional data from 35 interleaved slices (slice thickness=3mm; multi-slice mode=interleaved; FoV=192×192×105mm; voxel size=3mm isotropic; TR=2000ms, TE=30ms; flip-angle=90°; GRAPPA acceleration factor=2, time points=200, approximate acquisition time=7 minutes). Whole-brain T1-weighted anatomical images were acquired using a 3D MPRAGE sequence (slice thickness=0.9mm; FoV=263×350×350mm; voxel size=1mm isotropic; TR=1930ms; TE=2.36ms, flip-angle=8 °; GRAPPA acceleration factor=3).

#### ADNI

For ADNI, data was obtained using 3T MRI scanners with a standardised protocol across imaging sites (full details in https://adni.loni.usc.edu/wp-content/uploads/2017/07/ADNI3-MRI-protocols.pdf). An EPI-BOLD sequence was used to acquire functional data (slice thickness=3.4 mm, FoV=220×220×163mm, voxel size=3.4 isotropic; TR=3000ms; TE=30ms; flip angle=90°; GRAPPA acceleration factor=2; time points =197, approximate acquisition time=10 minutes). Whole-brain T1-weighted anatomical images were acquired with a 3D MPRAGE sequence (slice thickness=1 mm, FoV=208×240×256mm; voxel size=1×1×1 mm; TR=2300ms, TE=3ms, flip angle=9°, GRAPPA acceleration factor=3).

### Image processing

Image processing pipelines for the two cohorts included substantially similar steps, yet with some small differences (details below). Results are therefore not only replicated across different cohorts, but also irrespective of minor differences in preprocessing choices.

#### Geneva

fMRI data were preprocessed using in-house MATLAB code based on state-of-the-art fMRI processing guidelines ^42,74,82^. Below follows a brief description of these steps. Structural images were first denoised to improve the signal-to-noise ratio ^83^, bias-field corrected, and then segmented (FSL FAST) to extract white matter, grey matter and cerebrospinal fluid (CSF) tissue masks. These masks were warped in each individual subject’s functional space by means of subsequent linear and non-linear registrations (FSL flirt 6dof, FSL flirt 12dof and FSL fnirt). The following steps were then applied on the fMRI data: BOLD volume unwarping with applytopup, slice timing correction (slicetimer), realignment (mcflirt), normalisation to mode 1000, demeaning and linear detrending (MATLAB detrend), regression (MATLAB regress) of 18 signals: 3 translations, 3 rotations, and 3 tissue-based regressors (mean signal of whole-brain, white matter (WM) and cerebrospinal fluid (CSF), as well as 9 corresponding derivatives (backwards difference; MATLAB). We tagged high head motion volumes on the basis of two metrics: frame displacement (FD, in mm), and DVARS (D referring to temporal derivative of BOLD time courses, VARS referring to root mean square variance over voxels) as in 42. Specifically, we used the standardised DVARS as proposed in 88. We also used SD (standard deviation of the BOLD signal within brain voxels at every time-point). The FD and DVARS vectors (obtained with fsl_motion_outliers) were used to tag outlier BOLD volumes with FD > 0.3 mm and standardised DVARS > 1.7. The SD vector obtained with MATLAB was used to tag outlier BOLD volumes higher than the 75th percentile +1.5 of the interquartile range as per FSL recommendation ^84^. Subjects (N=4) with more than 30% motion-tagged volumes were excluded from the analyses. For the remaining subjects, the tagged volumes were not removed. There was no significant difference across groups in the percentage of tagged volumes [p>.928], while there was a tendency of higher FD in CU Aβ-relative to MCI Aβ+ and Dementia Aβ+ [p>.048]. There was no difference across test and retest in the percentage of tagged volumes [p=.471] nor in the average FD [p=.364]. Note that motion was accounted for in our statistical analyses (see section “Functional Connectivity and whole-brain within-session brain-fingerprint”).

A bandpass first-order Butterworth filter [0.01 Hz, 0.15 Hz] was applied to all BOLD time-series at the voxel level (MATLAB butter and filtfilt). The first three principal components of the BOLD signal in the WM and CSF tissue were regressed out of the grey matter (GM) signal (MATLAB, pca and regress) at the voxel level. A whole-brain data-driven functional parcellation based on 248 regions including cortical and subcortical areas as obtained by ^85^, was projected into each subject’s T1 space (FSL flirt 6dof, FSL flirt 12dof and finally FSL fnirt) and then into the native EPI space of each subject. We also applied FSL boundary-based-registration ^86^ to improve the registration of the structural masks and the parcellation to the functional volumes.

In some rare cases, BOLD signal from some ROIs was missing. When signal from an ROI was not available in more than 10% of subjects it was excluded from the analyses for all; this concerned a total of 8 ROIs, corresponding to 7 subcortical and 1 cortical ROIs. In the remaining few cases with no signal, metrics were computed with available data only.

#### ADNI

Anatomical and functional images were preprocessed with a standardised in-house-developed preprocessing pipeline ^87^ implemented in MATLAB (MATLAB 2021a version 9.10; MathWorks Inc., Natick, MA, USA) and including functions from SPM8 and SPM12 (http://www.fil.io-n.ucl.ac.uk/spm/). Individual structural T1 images were registered to each individual’s functional space (SPM coreg) while keeping the high T1 resolution, and segmented into grey matter, white matter and cerebrospinal fluid (SPM New Segment). Functional scans were realigned (SPM realign) and spatially smoothed (SPM smooth) with a Gaussian filter with FWHM=5mm. Nuisance signals were regressed out by means of a GLM, specifically linear and quadratic trends, 6 motion parameters and average signals in the white matter and cerebrospinal fluid. The same whole-brain data-driven functional parcellation ^85^ used for the Geneva dataset was adopted here to extract regional timecourses. The parcellation in MNI coordinates was first normalised to the individuals’ previously registered T1 images (functional space, structural high resolution), and then resampled to the lower functional BOLD resolution. Regional time series were then extracted by averaging the voxelwise preprocessed BOLD signals within each of the 248 regions of the parcellation. Finally, regional timecourses were band-pass filtered with cut-offs of 0.01-0.15 Hz to isolate typical resting-state fluctuations.

The FD vectors (obtained from SPM head motion parameters using the procedure described in ^74^) were used to tag outlier BOLD volumes with FD > 0.5 mm as per recommendation in ^75^. Subjects (N=7) with more than 30% motion-tagged volumes were excluded from the analyses (see section 1.0). For the remaining subjects, the tagged volumes were not removed. There was no significant difference across groups neither in the percentage of tagged volumes [p>=.634] nor in the average FD [p>=.897], while there was a difference across test and retest in the percentage of tagged volumes [p=.026] and in the average FD [p<.001], revealing that subjects moved more in the second part of the acquisition. To factor out the effect of motion in the brain-fingerprint, motion was added as nuisance variable in the whole brain-fingerprinting analyses (cf. section “Whole-brain within-sessions brain fingerprint during cognitive decline”).

In some rare cases, the BOLD signal from some ROIs was missing. When signal from an ROI was not available in more than 10% of subjects it was excluded from the analyses for all; this concerned a total of 2 ROIs in subcortical regions. In the remaining few cases with no signal, metrics were computed with available data-only.

### Functional Connectivity and whole-brain within-session brain-fingerprint

We estimated individual FC matrices using Pearson’s correlation coefficient between the averaged signals of all region pairs. The resulting individual FC matrices were composed of 248 nodes, as obtained by ^85^. Finally, the resulting functional connectomes were ordered according to seven cortical resting state networks (RSNs) as proposed by ^45^, plus one additional network including subcortical regions (similarly to ^88^, see also Fig. 1A).

We estimated *within-session* identifiability or fingerprinting, following the approach proposed in ^12^. This method involves splitting the fMRI times series in halves and enables quantification of within*-session connectome fingerprints.* Previous work has demonstrated that this method produces very similar results to those obtained from data acquired across separate sessions (i.e., between-sessions fingerprint) in healthy subjects from the Human Connectome Project (HPC) (see Fig. S3 in ^12^). Although *within* and *between-sessions fingerprinting* held similar results, they are different approaches to quantifying the brain-fingerprint, and this should be compared in future studies. However, we note that there are currently no clinical datasets available that include two fMRI sessions acquired within a short-time gap (i.e., within around one or two weeks). Therefore, *within-session fingerprint* is currently the only method available for estimating brain-fingerprint during cognitive decline. In this current study, we estimated identifiability across the first half (test) and second half volumes (retest) within the same scan. Recent work has shown that a good level of identifiability across the different resting state networks can be reached from around 200s (see Fig 4B, in ^75^). In this work, each test and retest session had 100 volumes (Geneva) or 90 volumes (ADNI) with a TR of 2s and 3s respectively, therefore providing sufficient data for achieving a good success rate and identifiability across the entire brain.

At the whole-brain level, the fingerprint was calculated for each subject *s* as test-retest similarity between FCs (cf. Fig 1B; we called this metric *ISelf*).

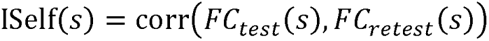

Then, for each subject *s* we computed an index of the FCs similarity with the other subjects *i* in their group (*IOthers*), where *N* is the total number of subjects in each group:

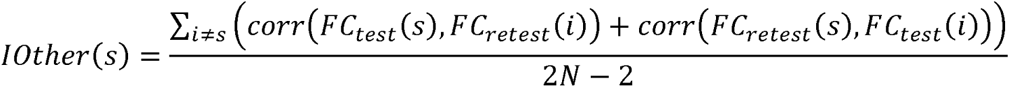

A second metric, *IDiff* (Fig. 1B), provides a group-level estimate of the within-(*ISelf*) and between-subjects (*IOthers*) test-retest reliability distance, where *subj* is the set of subjects:

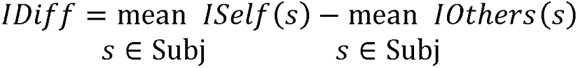

Finally, we measured the Success-rate ^7^ of the identification procedure as the percentage of cases with higher within-(*ISelf*) vs. between-subjects (*IOthers*) test-retest reliability. These metrics have been introduced and estimated in healthy populations in previous work ^12^. We performed paired-sample t-test to compare *ISelf* vs. *IOthers* in each group/cohort. Then, we used one-way ANOVAs to test the effect of group on *ISelf* and *IOthers* separately after checking for nuisance variables, with 5000 permutations to control for sample size differences. For *ISelf*, the nuisance variables were age, sex, years of education (YoE), and the difference in motion between the test and retest scans, as absolute difference between FD. For *IOthers*, the nuisance variables were also age, sex, and YoE, but motion (FD) was considered across the entire acquisition, as *IOthers* is a composite measure across test and retest. Additionally, when the scanner type varied across subjects (i.e., in ADNI), scanner type was also included as a nuisance variable for *IOthers*. Finally, we did a permutation testing analysis to compare *Success-rate* and *IDiff* from 1000 surrogate datasets of random ID matrices against the real value ^46^.

### Spatial specificity of brain fingerprint: edge-wise intra-class correlation

Spatial specificity of FC fingerprints was derived using edgewise intra-class correlation (ICC) with one-way random effect model according to ^64^ (cf. Fig. 1C):

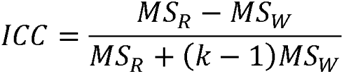

Where *MS_R_* = mean square for rows (between the subjects); *MS_W_* = mean square for residual sources of variance; *k* = sessions. ICC coefficients quantify the degree of similarity between observations/measures and find high applicability in reliability studies ^89^. The higher the ICC coefficient, the stronger the agreement between two observations. Here we used this metric, as in previous work ^12,75^, to quantify the similarity between test and retest for each edge (FC between 2 regions). A high ICC indicates that a larger proportion of the variance across test and retest is due to differences between the subjects, rather than differences between test and retest or random error. A low ICC, on the other hand, indicates that there is more variability due to differences between test and retest or random error, than due to differences between subjects. In other words, the higher the ICC of an edge, the more that edge connectivity is similar for each subject across test and retest, as well as the variability across subjects, i.e., the higher the ‘fingerprint’ of that edge.

Edge-wise ICC was computed for all possible edges and for each group separately, with the aim to quantify the edges-wise functional connectivity fingerprint, distinctive of each clinical group. In order to control for sample size differences across groups, bootstrapping was used to accurately estimate edgewise fingerprints: for each group, ICC was calculated across test and retest for subsets of randomly chosen N=10 subjects, across 1000 bootstrap runs, and then averaged within each group (Fig. 3A and 3B). Matrices in 3A and 3B were binarized for ICC>0.6 which is considered the lower threshold for a ‘good’ ICC score ^47^, and overlapping edges across the two cohorts were displayed in the binary ICC matrices (cf. overlap ICC matrix in Fig. 3C).

### Brain fingerprint in resting-state functional networks during cognitive decline

Next, we aimed to identify the commonalities in the distribution of edges with the highest ICC across different cohorts, both within and between resting-state networks. To achieve this, we analysed the overlap ICC binary matrix (Fig. 3C) and computed the following metrics. For each of one seven resting state networks, both within and between-networks (*net*), we quantified the number of overlapping ICC edges (*ICC_O_*). Next, we computed a proportion of the *ICC_O_* edges relative to the total number of edges in each network and defined it as 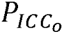. Finally, and in order to determine the distance from the healthy reference (i.e., CU), we computed the ratio over healthy individuals (Fig. 4A). The ratio (R) was calculated as follow, where *D* = disease (i.e., MCI and Dementia) and *H* = healthy (i.e., CU).

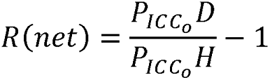

Lastly, we aimed to determine whether there was a significant difference in the distribution of overlapping edges in within-networks vs. between-networks for each group. To achieve this, we conducted a Chi-square test for each network comparing the number of edges in the within-versus the between-functional networks. Significance was Bonferroni corrected for multiple comparisons (Fig. 4B).

### Between-groups significance of nodal brain fingerprint

In these analyses, we aimed to identify regions (or nodes) whose functional connectivity with the rest of the brain could account for significant differences in subject variability across the three groups. First, to isolate edges that were significantly different across groups, we compared real ICC matrices with surrogate ones obtained after including randomly selected subjects from each group for 1000 permutation runs (Fig. S3), i.e., we generated surrogate *group-unspecific* ICC matrices. Next, a p-value was computed for each edge as a proportion of times across permutation runs where surrogate values were bigger than the real value; edges with p<.05 were considered significant (see Fig. S3). Only significant edges were selected and a new matrix including significant edges with its real ICC value and having zeros for all the non-significant ones was used to compute nodal strength ICC. Nodal strength was computed as average, including zeros to account for non-significant edges, and rendered on the cortical surface using BrainNet ^90^ (cf. Fig. S4). Next, to select the group-specific nodes with the highest ICC values that were common across the two independent cohorts, binary masks were obtained by selecting the top 25 percentile of ICC nodal strength and the overlap between the two cohorts was displayed (Fig. 5A). Each binary mask obtained that way provides a nodal representation of the brain region “hubs” involved in FC fingerprints in each group specifically.

### Brain fingerprint across cognitive decline and behaviour

A Neurosynth meta-analysis (https://neurosynth.org/), similar to the one implemented in previous studies ^53,54^, was conducted to assess cognitive functions associated with brain fingerprints at the different stages of cognitive decline. The procedure outputs, for each combination of brain fingerprint mask and cognitive function binary mask, a nodal z statistic that quantifies the similarity between the two maps. For brain fingerprint, we used the binary overlap masks in Fig. 5A – i.e., those including the ICC hubs with the highest fingerprint across the two cohorts. For cognition, we used the brain binary maps related to 50 topic terms common in the neuroimaging literature ^73,91^ derived from the Neurosynth database. These fingerprint and cognition maps were used as input for the meta-analysis to find significant associations between the ICC hub or fingerprint masks and the brain cognitive functions Neurosynth maps. Last, we ordered the terms according to the weighted mean of the resulting z statistics for visualization, considering significant any association between group fingerprints and cognitive maps above z>3.1 ^53,54^ (Fig. 5B).

## Supporting information

Supplementary Materials, revised

## Acknowledgements

SS was supported by grants from the Swiss National Science Foundation [SNSF 320030_185028, PI: VG]. Data acquisition from the Geneva cohort was granted by SNSF 320030_185028 and 169876 (PI: VG) and Fonds Startup du département de radiologie et informatique médicale, Université de Genève, Faculté de Médecine (PI: SS).

The Geneva Memory Centre (Centre de la Mémoire) is funded by the following private donors under the supervision of the Private Foundation of Geneva University Hospitals: A.P.R.A. - Association Suisse pour la Recherche sur la Maladie d’Alzheimer, Genève; Fondation Segré, Genève; Race Against Dementia Foundation, London, UK; Fondation Child Care, Genève; Fondation Edmond J. Safra, Genève; Fondation Minkoff, Genève; Fondazione Agusta, Lugano; McCall Macbain Foundation, Canada; Nicole et René Keller, Genève; Fondation AETAS, Genève. Competitive research projects have been funded by: H2020 (projects n. 667375), Innovative Medicines Initiative (IMI contract n. 115736 and 115952), IMI2, Swiss National Science Foundation (projects n.320030_182772 and n. 320030_169876), VELUX Foundation. The Clinical Research Center, at Geneva University Hospital and Faculty of Medicine provides valuable support for regulatory submissions and data management. The authors thank Avid Radiopharmaceuticals Inc. for providing the 18F-Flortaucipir tracer without being involved in the data analysis or interpretation.

EA acknowledges financial support from the SNSF Ambizione project “Fingerprinting the brain: Network science to extract features of cognition, behaviour and dysfunction” (grant number PZ00P2_185716). VG is supported also from the Velux Foundation, the Aetas Foundation, the Schmidheiny Foundation and research/teaching support through her institution from Siemens Healthineers, GE Healthcare, Novo Nordisk and Janssen. MP is partially supported by the Italian Ministry of Health (Ricerca Corrente). MGP was supported by the CIBM Center for Biomedical Imaging, a Swiss research center of excellence founded and supported by Lausanne University Hospital (CHUV), University of Lausanne (UNIL), Ecole polytechnique fédérale de Lausanne (EPFL), University of Geneva (UNIGE) and Geneva University Hospitals (HUG).

Data collection and sharing for the ADNI cohort was funded by ADNI (National Institutes of Health Grant U01 AG024904) and DOD ADNI (Department of Defense award number W81XWH-12-2-0012). ADNI is funded by the National Institute on Aging, the National Institute of Biomedical Imaging and Bioengineering, and through generous contributions from the following: AbbVie, Alzheimer’s Association; Alzheimer’s Drug Discovery Foundation; Araclon Biotech; BioClinica, Inc.; Biogen; Bristol-Myers Squibb Company; CereSpir, Inc.; Cogstate; Eisai Inc.; Elan Pharmaceuticals, Inc.; Eli Lilly and Company; EuroImmun; F. Hoffmann-La Roche Ltd and its affiliated company Genentech, Inc.; Fujirebio; GE Healthcare; IXICO Ltd.; Janssen Alzheimer Immunotherapy Research & Development, LLC.; Johnson & Johnson Pharmaceutical Research & Development LLC.; Lumosity; Lundbeck; Merck & Co., Inc.; Meso Scale Diagnostics, LLC.; NeuroRx Research; Neurotrack Technologies; Novartis Pharmaceuticals Corporation; Pfizer Inc.; Piramal Imaging; Servier; Takeda Pharmaceutical Company; and Transition Therapeutics. The Canadian Institutes of Health Research is providing funds to support ADNI clinical sites in Canada. Private sector contributions are facilitated by the Foundation for the National Institutes of Health (www.fnih.org). The grantee organization is the Northern California Institute for Research and Education, and the study is coordinated by the Alzheimer’s Therapeutic Research Institute at the University of Southern California. ADNI data are disseminated by the Laboratory for Neuro Imaging at the University of Southern California.

We thank Benedetta Franceschiello, Emahnuel Troisi-Lopez, Pierpaolo Sorrentino, Valerio Zerbi and Jenya Chumin for the insightful comments. We are indebted to the patients, their carers, and the volunteers for taking part in this study, and with the clinical and research staff of the Geneva Memory Centre for data acquisition.

## Authors contributions

Authors contribution according to the CRediT taxonomy, see http://credit.niso.org/ for more information.

*Conceptualisation*: SS and EA (lead); MP, DVDV, GBF, OB and VG contributed to the evolution of the overarching research goals and aims.

*Methodology*: SS and EA (lead), DVDV contributed to the creation of analytical models;

*Software*: SS and EA wrote the code for the analyses;

*Validation*: SS;

*Formal analyses*: SS (lead), EA, SA, AF and MGP contributed to fMRI data preprocessing;

*Resources:* MS, KOL, AF, MGP, PU, GBF, VG, EA;

*Data curation:* SS (lead), ST, FR, AF, VG, MGP contributed to data curation;

*Writing – Original Draft*: SS (lead) and EA;

*Writing – Review & Editing:* SS, EA, VG, OB, DVDV, MS, FR, MGP, MP;

*Visualisation:* SS (lead) and EA;

*Supervision*: EA, VG;

*Project Administration:* SS, EA, VG;

*Funding acquisition:* SS, EA, VG, OB, GBF.

