## Supplementary Materials, revised for "Fingerprints of brain disease: Connectome identifiability in cognitive decline and Alzheimer’s disease"

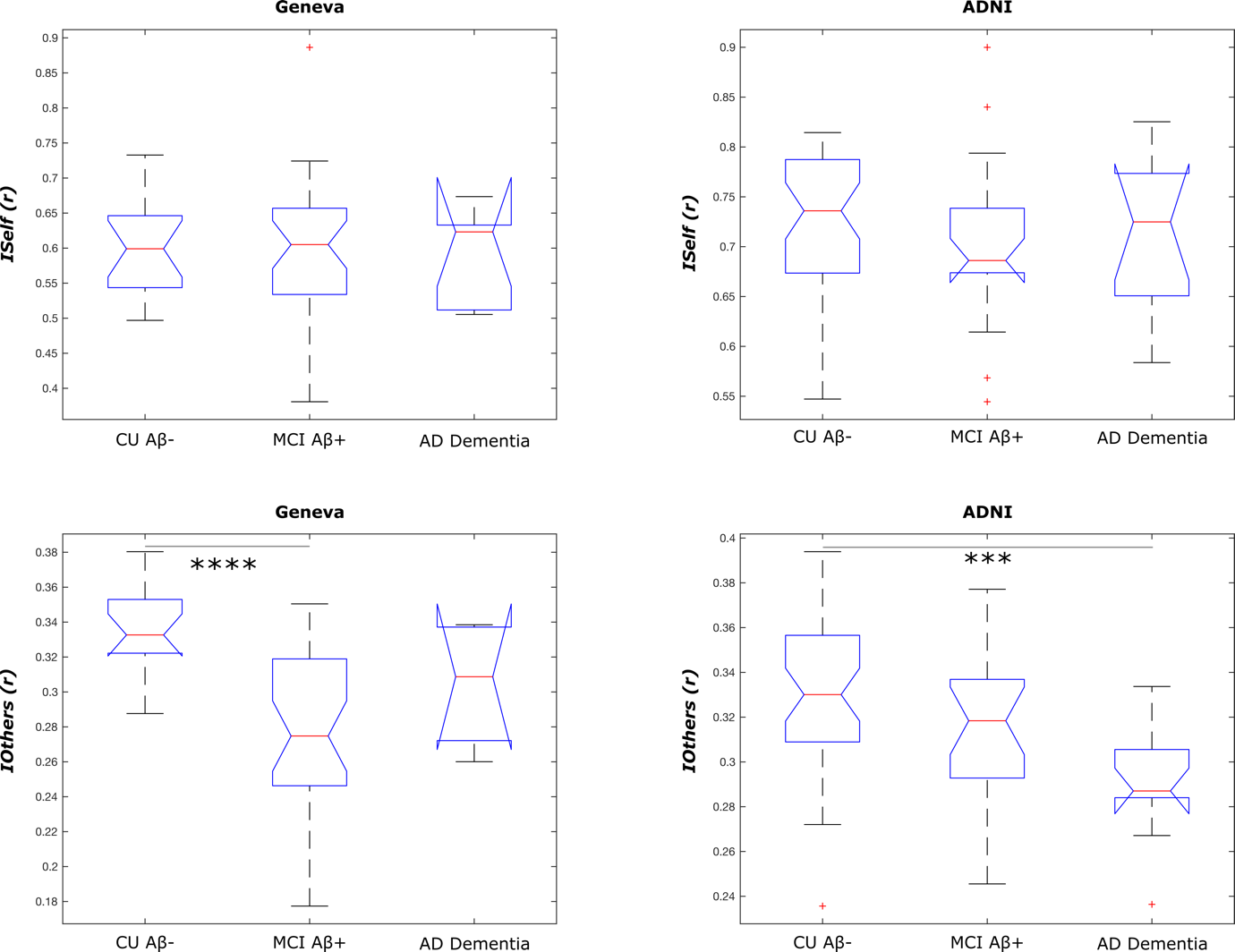

**Supplementary Figure 1.** **ISelf (A) and IOthers (B) across groups.** **A)** Boxplots show that, both in Geneva and ADNI cohort, *ISelf* was equivalent in the three groups. **B)** *IOthers* differed across the three groups in both Geneva and ADNI cohorts. Relative to the healthy reference (CU Aβ-), MCI Aβ+ and AD Dementia patients had lower *IOthers* in Geneva and ADNI, respectively. **** p ≤ 0.0001; *** p ≤ 0.001. Significance assessed through post-hoc pairwise comparisons with Bonferroni correction. Absence of * denote no significant difference across groups. Cf. Supplementary Tables 1 for full statistics on one-way ANOVA and subsequent post-hoc pairwise comparisons.

Supplementary Table 1.

A) Effect of group on *ISelf*: One-way ANOVA results – Geneva and ADNI.

| *Geneva* | | | | | | |
| --- | --- | --- | --- | --- | --- | --- |
| *Anova Table* | SS | df | F | parametric p | resampled p | p resampled signif. |
| Group | 0.0013 | 2 | 0.08 | 0.921 | 0.920 | ns |
| Age | 0.0010 | 1 | 0.13 | 0.721 | 0.713 | ns |
| Sex assigned at birth | 0.0175 | 1 | 2.21 | 0.145 | 0.141 | ns |
| YoE | 0.0066 | 1 | 0.84 | 0.364 | 0.378 | ns |
| Δ FD | 0.0009 | 1 | 0.12 | 0.729 | 0.734 | ns |
| *ADNI* | | | | | | |
| *Anova Table* | SS | df | F | parametric p | resampled p | p resampled signif. |
| Group | 0.0095 | 2 | 0.95 | 0.394 | 0.389 | ns |
| Age | 0.0014 | 1 | 0.28 | 0.598 | 0.602 | ns |
| Sex | 0.0185 | 1 | 3.66 | 0.060 | 0.058 | ns |
| YoE | 0.0027 | 1 | 0.54 | 0.464 | 0.465 | ns |
| Δ FD | 0.0143 | 1 | 2.84 | 0.097 | 0.096 | ns |

*Legend:* One-way ANOVAs to test the effect of group on IOthers after checking for nuisance variables (Age, Sex, YoE and absolute difference between motion (Δ FD) at test vs. retest), with 5000 permutations to control for sample size differences. Resampling test using Freedman-Lane to handle nuisance variables and 5000 permutations; YoE=Years of Education; Δ FD= absolute difference between test and retest in Frame displacement; ns=p>0.05.

B) Effect of group on *IOthers*: One-way ANOVA results – Geneva and ADNI

| *Geneva* | | | | | | |
| --- | --- | --- | --- | --- | --- | --- |
| *Anova Table* | SS | df | F | parametric p | resampled p | p resampled signif. |
| Group | 0.0289 | 2 | 15.27 | 0.000 | 0.000 | *** |
| Age | 0.0001 | 1 | 0.14 | 0.715 | 0.716 | ns |
| Sex assigned at birth | 0.0006 | 1 | 0.63 | 0.432 | 0.434 | ns |
| YoE | 0.0004 | 1 | 0.39 | 0.535 | 0.542 | ns |
| FD | 0.0219 | 1 | 23.17 | 0.000 | 0.000 | *** |
| *ADNI* | | | | | | |
| *Anova Table* | SS | df | F | parametric p | resampled p | p resampled signif |
| Group | 0.0101 | 2 | 4.95 | 0.010 | 0.009 | ** |
| Age | 0.0022 | 1 | 2.19 | 0.144 | 0.133 | ns |
| Sex assigned at birth | 0.0052 | 1 | 5.14 | 0.027 | 0.026 | * |
| YoE | 0.0010 | 1 | 0.96 | 0.332 | 0.334 | ns |
| FD | 0.0005 | 1 | 0.45 | 0.506 | 0.529 | ns |
| Scanner | 0.0063 | 6 | 1.02 | 0.418 | 0.420 | ns |

*Legend:* One-way ANOVAs to test the effect of group on IOthers after checking for nuisance variables (Age, Sex, YoE and absolute difference between motion (FD) at test vs. retest and Scanner model), with 5000 permutations to control for sample size differences. Resampling test using Freedman-Lane to handle nuisance variables and 5000 permutations; YoE=Years of Education; FD= Frame displacement. ns = p > 0.05; * p ≤0.05, ** p ≤0.01, *** p ≤ 0.001.

C) Effect of group on *IOthers*: post-hoc pairwise comparisons – Geneva and ADNI.

| *Geneva* | | | | | | | | |
| --- | --- | --- | --- | --- | --- | --- | --- | --- |
| term | y | group1 | group2 | df | statistic | p | p adj. | p adj. signif |
| FD*Group | Iothers | CU | MCI | 50 | 2.34 | 0.023 | 0.069 | ns |
| FD*Group | Iothers | CU | Dementia | 50 | 5.90 | 0.000 | 0.000 | **** |
| FD*Group | Iothers | MCI | Dementia | 50 | 1.53 | 0.132 | 0.397 | ns |
| *ADNI* | | | | | | | | |
| term | y | group1 | group2 | df | statistic | p | p adj. | p adj.  signif |
| Group | Iothers | CU | MCI | 69 | -3.23 | 0.002 | 0.006 | *** |
| Group | Iothers | CU | Dementia | 69 | -1.77 | 0.081 | 0.244 | ns |
| Group | Iothers | MCI | Dementia | 69 | 1.64 | 0.107 | 0.320 | ns |

*Legend:* Pairwise comparisons between groups using the estimated marginal means. p adj. = p value adjusted using Bonferroni correction for multiple comparisons. ns = p > 0.05; *** p ≤ 0.001; **** p ≤ 0.0001.

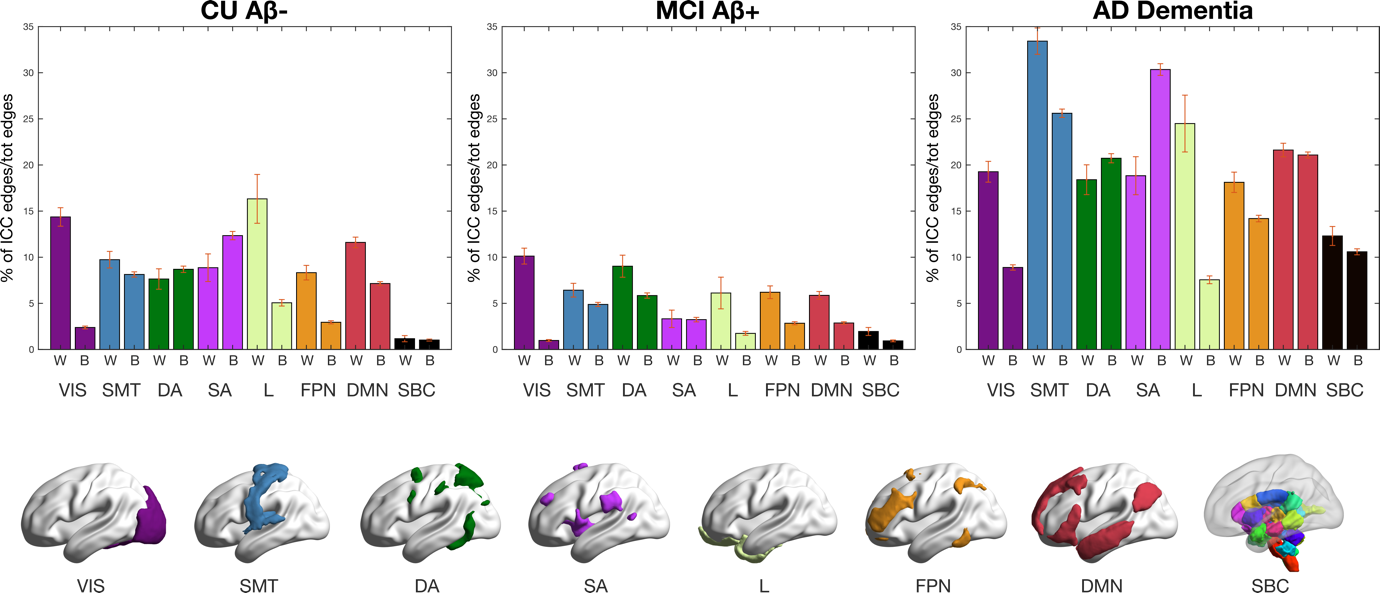

**Supplementary Figure 2. Distribution of brain fingerprint across resting-state functional networks.** Distribution of the edges with highest ICC common to both cohorts (from ICC overlap binary matrix, cf. Fig. 3C) in within-networks and between-networks. We display percentages of edges out of the total number of edges for each within-network and between-network. Error bars represent standard error. W=within-networks; B=between-networks; ICC=intra-class correlation; CU: Cognitively Unimpaired; MCI: Mild Cognitive Impairment. VIS=visual network; SMT=somatomotor network; DA=dorsal-attention network; SA=salience network; L=limbic network; FPN=fronto-parietal network; DMN=default-mode network; SBC=subcortical regions.

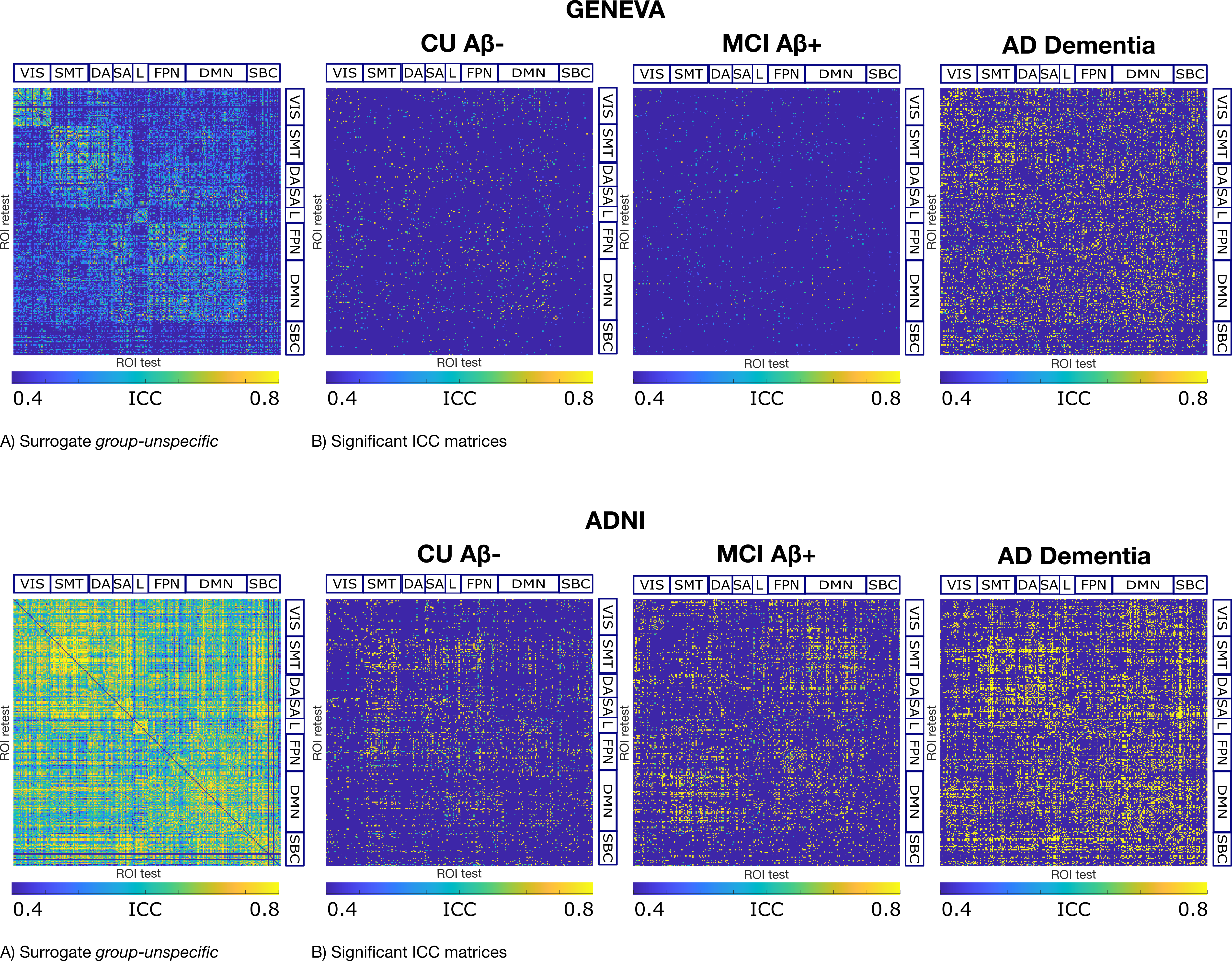

**Supplementary Figure 3. A)** Surrogate group-unspecific ICC matrix, obtained after including randomly selected subjects from each group for 1000 permutation runs. **B)** ICC matrix including significant edges only; a p-value was computed for each as a proportion of times across permutation runs where surrogate values were bigger than the real value; edges with p <.05 were considered significant and displayed here. Geneva and ADNI cohorts in first and second row, respectively. ICC=intra-class correlation; CU: Cognitively Unimpaired; MCI: Mild Cognitive Impairment. VIS=visual network; SMT=somatomotor network; DA=dorsal-attention network; SA=salience network; L=limbic network; FPN=fronto-parietal network; DMN=default-mode network; SBC=subcortical regions.

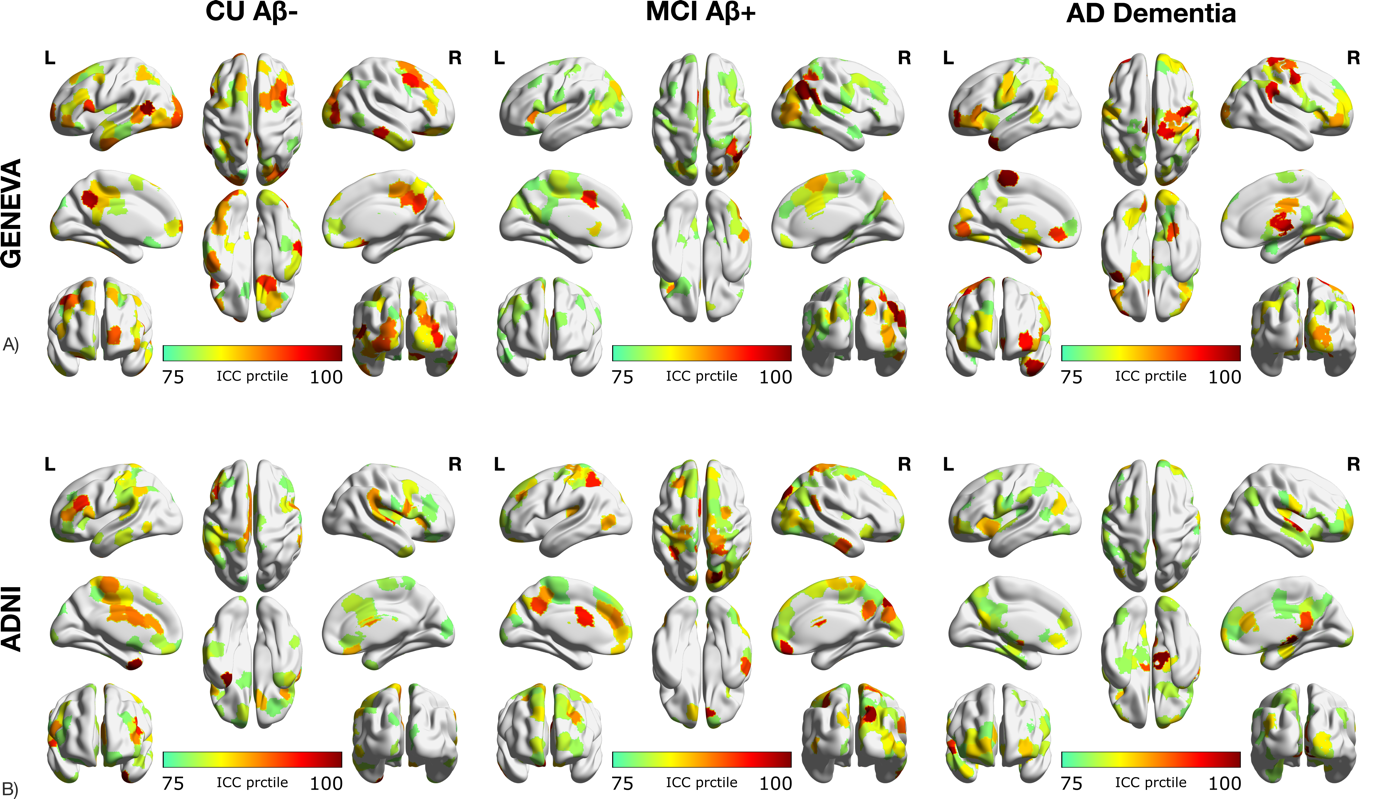

**Supplementary Figure 4.** Brain renders showing nodal strength for each group in **A)** Geneva and **B)** ADNI. Nodal strength was computed from significant ICC matrices (cf. S. Fig. 3) as average, including zeros to account for non-significant edges, and rendered on the cortical surface using BrainNet^1^. Only nodal strength values above the 75th percentile were selected.
